# AtDAT1 is a key enzyme of D-amino acid stimulated ethylene production in *Arabidopsis thaliana*

**DOI:** 10.1101/716373

**Authors:** Juan Suarez, Claudia Hener, Vivien-Alisa Lehnhardt, Sabine Hummel, Mark Stahl, Üner Kolukisaoglu

## Abstract

D-enantiomers of proteinogenic amino acids (D-AAs) are found ubiquitously, but the knowledge about their metabolism and functions in plants is scarce. A long forgotten phenomenon in this regard is the D-AA-stimulated ethylene production in plants. As a starting point to investigate this effect the *Arabidopsis* accession Landsberg erecta (Ler) got into focus as it was found defective in metabolizing D-AAs. Combining genetics and molecular biology of T-DNA lines and natural variants together with biochemical and physiological approaches we could identify AtDAT1 as a major D-AA transaminase in *Arabidopsis. Atdat1* loss-of-function mutants and *Arabidopsis* accessions with defective *AtDAT1* alleles were not able to produce D-Ala, D-Glu and L-Met, the metabolites of D-Met, anymore. This result corroborates the biochemical characterization of AtDAT1, which showed highest activity using D-Met as substrate. Germination of seedlings in light and dark led to enhanced growth inhibition of *atdat1* mutants on D-Met. Ethylene measurements revealed an enhanced D-AA stimulated ethylene production in these mutants. According to initial working models of this phenomenon D-Met is preferentially malonylated instead of the ethylene precursor 1-aminocyclopropane-1-carboxylic acid (ACC). This decrease of ACC degradation should then lead to the increase of ethylene production. We could observe in our studies a reciprocal relation of malonylated methionine and ACC upon D-Met application and even significantly more malonyl-methionine in *atdat1* mutants. Unexpectedly, the malonyl-ACC levels did not differ between mutants and wild type in these experiments. With AtDAT1, the first central enzyme of plant D-AA metabolism was characterized biochemically and physiologically. The specific effects of D-Met on ACC metabolization, ethylene production and plant development of *dat1* mutants unraveled the impact of AtDAT1 on these processes, but they are not in full accordance to previous working models. Instead, our results imply the influence of additional candidate factors or processes on D-AA-stimulated ethylene production which await to be uncovered.

## 1 Introduction

It is widely accepted that proteinogenic L-amino acids (L-AAs) are essential in all kingdoms of life, both as primary metabolites as well as elementary building blocks of proteins. In contrast, the metabolism and functions of the D-forms of amino acids (D-AAs) is far less clear and defined. Major reasons for this discrepancy are the large diversity and different functions of D-AAs in organisms. For instance, bioactive peptides like octopine from octopus and scallop, antibiotics from bacteria and opioids from frogs were among the first substances reported to contain D-AAs. (Fujii, 2002;Martínez-Rodríguez et al., 2010;Ollivaux et al., 2014) In humans, several proteins related to diseases like arteriosclerosis, Alzheimer or Parkinson contain D-AAs, especially D-Asp, which are generated by racemization of the corresponding L-AA (Fujii et al., 2011). Various free D-AAs were detected in different tissues and fluids of humans and other mammals (Hamase et al., 2002;Hamase, 2007). The most prominent example in this respect is the impact of D-Asp and D-Ser on the functions of the N-methyl-D-aspartate (NMDA) receptor in mammals: Aberrant levels of these D-AAs seem also to be connected with psychological disorders and diseases of the endocrine system (for reviews see Fuchs et al. (2005);D’Aniello (2007);Katane and Homma (2011);Balu and Coyle (2015)).

Far less is known about the metabolism and functions of D-AAs in plants. This is astonishing against the background that plant roots are surrounded by D-AAs, most obvious, but not exclusively D-Ala and D-Glu, as degradation products of the peptidoglycan layer of bacterial cell walls (Dworkin, 2014). Thus, the amount of D-AAs in the rhizosphere can be more than 10% of the corresponding L-enantiomer (Brodowski et al., 2005;Amelung et al., 2006). This led to the question if D-AAs are actively utilized by plants. For a long time, D-AAs were considered as toxins due to the fact, that some of them inhibit seedling growth in submillimolar concentrations (Erikson et al., 2004;Forsum et al., 2008). But several reports suggest that D-AAs take up a similarly crucial position in plants as in microbes and animals (for further readings about D-AAs in microbes and animals see Konno et al. (2007), and Brückner (2011)). For instance, the D-Ala amount in duckweed (*Landoltia punctata*) was demonstrated to be increased during UV stress (Monselise et al., 2015) and D-Ser is involved in pollen tube growth in *Arabidopsis* by regulating the glutamate receptor GLR1.2, which belongs to a group of plant proteins closely related to mammalian NMDA receptors (Michard et al., 2011;Forde and Roberts, 2014). In mosses (*Physcomitrella patens*) D-Ala and D-Glu were detected in the plastidial envelope, which resembles to bacterial peptidoglycan (Hirano et al., 2016). This finding and others led to the conclusion that peptidoglycan, containing D-Ala and D-Glu, is an integral part of the plastidial envelope not only in cryptophytes (for a review see Chen et al. (2018)).

The number of enzymes predicted to be specific for processing D-AAs annotated in plant genomes imply much more functions for these amino acids than currently known (Naranjo-Ortíz et al., 2016). However, it also raises the question about its metabolism in plants, especially how the abundance of different D-AAs is regulated. On the one hand their contents have to be maintained at required levels to ensure their activity. On the other hand, the intracellular concentrations must be limited below toxic levels. This restriction is of specific importance due to the fact that the rhizosphere is the natural major source of D-AAs for plants (Vranova et al., 2012) and that D-AAs are taken up by roots in considerable amounts compared to L-AAs (Hill et al., 2011;Gördes et al., 2013). In this respect, the question arises which processes facilitate the catabolization of D-AAs in plants.

In the course of these previous studies D-Met got into our focus. This D-AA caused the greatest conversion rates in almost all tested accessions of *Arabidopsis thaliana* except in Ler (Gördes et al., 2013). This attracted our interest insofar as methionine represents a relatively small portion of soil amino acids (Vranova et al., 2012). But it had been detected in soil (Amelung and Zhang, 2001), and there have also been several bacterial species isolated from soil which are specialized to the utilization of D-Met as sole carbon and nitrogen source (Radkov et al., 2016). Furthermore, it is produced by different bacteria, incorporated into their cell wall and even released to their environment in order to disassemble biofilms (for a review see Cava et al. (2011)). Nevertheless, D-Met has not been reported yet to be produced by plants.

More than 30 years ago it was reported that feeding D-Met to seedlings of cocklebur (*Xanthium pennsylvanicum*), pumpkin (*Cucurbita moschata*), sunflower (*Helianthus annuus*), mung bean (*Vigna radiata*), water melon (*Citrullus vulgaris*), and pea (*Pisum sativum*) leads to increased ethylene production (Satoh and Esashi, 1980;Liu et al., 1983;Kionka and Amrhein, 1984). This phenomenon was coined as “D-amino-acid-stimulated ethylene production” (Satoh and Esashi, 1980) due to the increase of ethylene content induced by D-Met and other D-AAs. The authors tried to explain the effect by competitive malonylation of D-Met and 1-aminocyclopropane-1-carboxylic acid (ACC), the precursor of ethylene. According to this hypothesis D-Met would compete with ACC for the same malonyl transferase (Liu et al., 1983;Ling-Yuan et al., 1985;Benichou et al., 1995;Wu et al., 1995), which would lead to an increase of ACC level and subsequently the ethylene production would rise (S F Yang and Hoffman, 1984). However, this hypothesis could not be verified because the corresponding malonyl transferase has not been identified to date. Nevertheless, the question about the mechanism and function of this phenomenon remains.

Our starting point to address this question were previous works from Gördes et al. (2011), which revealed that *Arabidopsis* plants convert particular D-AAs like D-Met, D-Trp, D-Phe and D-His partially to their respective L-enantiomers. Additionally, the feeding of almost all tested D-AAs led mainly to the formation of D-Ala and D-Glu. By contrast, the *Arabidopsis* accession Landsberg erecta (Ler) is incapable of both, the D-AA to L-AA and the D-AA to D-Ala/D-Glu conversion (Gördes et al., 2013). These observations point to a central metabolic step, in which D-AAs, with a high preference to D-Met, are converted to D-Ala and D-Glu by a D-AA specific transaminase (Vranova et al., 2012;Gördes et al., 2013).

In this paper we describe the identification and characterization of *Arabidopsis* loss-of-function mutant alleles in the Columbia-0 (Col-0) accession for a previously characterized D-AA specific transaminase called D-AAT (Funakoshi et al., 2008), which we named AtDAT1. Mutants of this gene showed almost identical defects as Ler to metabolize D-AAs, with D-Met as strongest effector. Indeed, we could show that the affected gene in Ler is an almost non-functional *AtDAT1* allele. Biochemical analyses revealed that this enzyme prefers D-Met as amino donor and pyruvate over 2-oxoglutarate as amino acceptor, confirming the preferential production of D-Ala in Col-0. The discovery of AtDAT1 and its mutants gave us also the opportunity to verify the working model of D-AA-stimulated ethylene production in plants. We found that D-Met causes significantly higher ethylene production and growth inhibition in *atdat1* seedlings. According to the current working model (see above) the increase in ethylene should be caused by a decrease in malonylation of ACC due to the increase of malonyl-D-Met, leading to higher ACC oxidation. Although we found higher malonyl-methionine concentrations in *atdat1* seedlings after D-Met feeding as expected, the malonyl-ACC levels decreased equally in mutants and their respective wild type. This points to an additional, yet unraveled, mechanism regulating D-AA-stimulated ethylene production in plants, which will be discussed. Furthermore, our findings point to functions of D-Met in central plant processes beyond unspecific growth inhibition.

## 2 Materials and Methods

### 2.1 Plant Material and Growth Conditions

All *Arabidopsis* ecotypes as well as T-DNA insertion lines analyzed in this study were either provided by the Nottingham Arabidopsis Stock Centre (University of Nottingham, UK) or the Arabidopsis Biological Resource Center (University of Ohio, Columbus, OH).

Seedlings for amino acid extraction and profiling were germinated in microtiter plates as described before (Gördes et al., 2013). For phenotypic analysis of seedlings and subsequent measurement of malonylated methionine and ACC in their extracts, plants were either germinated for six days in darkness or ten days in light (all at 22 °C). As solid growth media 1/2 MS basal salts with 1% sucrose and 1% phytoagar, including conditional further additions (e.g. D-AAs, ACC) were applied. For all analyses of adult plants these were grown in the greenhouse in soil.

### 2.2 PCR Genotyping and RT-PCR analysis of *Arabidopsis* Lines and Accessions

Plant DNA for PCR analysis was extracted from seedlings or leaves of adult plants according to Edwards et al. (1991). To determine zygosity of T-DNA insertion lines either a gene specific primer and a border primer or two gene specific primers flanking the insertion (for primer combinations and sequences see Supplemental Table S1) were used in a PCR reaction with Taq polymerase from New England Biolabs (Frankfurt am Main, Germany) according to manufacturer’s protocol. To determine the AtDAT1 sequence in different *Arabidopsis* ecotypes the complete coding sequences were amplified from genomic DNA and cDNA as described above and the PCR amplificates were sequenced directly by GATC (Konstanz, Germany). For cDNA synthesis RNA of 14 days old seedlings germinated in liquid media under long day conditions was extracted with the RNeasy Mini Kit from Qiagen (Düsseldorf, Germany) and cDNA was synthesized with RevertAid H Minus Reverse Transcriptase from Thermo Fisher Scientific (Karlsruhe, Germany), both according to manufacturers’ protocols. This cDNA was used for cloning purposes (see below) and RT-PCR analysis.

### 2.3 Cloning of AtDAT1 Variants for Recombinant Expression

For cloning AtDAT1 from cDNA of *Arabidopsis* accessions Col-0 and Ler, the complete coding sequence was amplified with KOD DNA Polymerase from Merck Millipore (Schwalbach am Taunus, Germany) with the primer combination DAT1-Start/DAT1-A1 (Supplemental Table S1). Amplificates were cloned into pENTR/D-TOPO according to manufacturer’s protocol (Thermo Fisher Scientific, Karlsruhe, Germany), leading to the constructs pENTR-AtDAT1_(Col-0)_ and pENTR-AtDAT1_(Ler)_. To create AtDAT1 coding sequences with the single point mutations A77T and T303S the previously described clones were cleaved with *Pst I* and *Not I*, creating a 0.5 kb fragment. This was then ligated from pENTR-AtDAT1_(Col-0)_ to pENTR-AtDAT1_(Ler)_ and vice versa, resulting in the constructs pENTR-AtDAT1_(A77T)_ and pENTR-AtDAT1_(T303S)_. After sequence verification of the constructs they were all used for LR reaction using the kit form Invitrogen (Karlsruhe, Germany) according to manufacturer’s protocol into pGEX-2TM-GW (kindly received from Bekir Ülker) for expression in *E. coli* with N-terminal GST tag and C-terminal His tag. Additionally, the pENTR-AtDAT1_(Col-0)_ and pENTR-AtDAT1_(Ler)_ were used for Gateway-based cloning into pUB-DEST-GFP for expression in plants with C-terminal GFP tag. pENTR-AtDAT1_(Col-0)_ was used for Gateway-based cloning into pUB-DEST (Grefen et al., 2010) for complementing AtDAT1 defective plants.

### 2.4 *Arabidopsis* Transformation and tobacco leaf infiltration

All plant transformation vectors were transformed into *Agrobacterium tumefaciens* cv. pMP90-RK GV3101. Plant transformation was performed by floral dipping (Clough and Bent, 1998). For selection of transformants, seeds were either germinated on 1/2 MS-Agar with 1% sucrose containing hygromycin or germinated on soil and sprayed with 2% BASTA from AgrEvo (Düsseldorf, Germany) depending on the used vector.

For tobacco leaf infiltration transformed *Agrobacterium* containing pUB10-GFP::DAT1 was mixed with a strain of transformed *Agrobacterium* for expression of the mCherry plastid marker (CD3-999 pt-rk; Nelson et al., 2007) and P19 *Agrobacterium tumefaciens* cells into infiltration media (10 mM MES-KOH [pH 5.7], 10 mM] MgCl_2,_ 0.2 mM Acetosyringone). Using a syringe 1mL of infiltration media with the mix of the 3 type of cells was infiltrated in the abaxial side of *Nicotiana benthamiana* leaves. Plants were then watered and kept on the lab bench for 2 d. Afterwards, single leaf discs were excised for confocal fluorescence microscopy.

### 2.5 Fluorescence Microscopy

Imaging was performed using a Leica laser scanning microscope SP8 with the corresponding software LCS or LASAF X (Leica Microsystems, Wetzlar, Germany). For excitation of GFP-fusion proteins the Argon laser was used at 488 nm and the detection range was from 500 to 550 nm. For m-RFP excitation was set to 561 nm and detection was from 600 to 650 nm. All autofluorescence of chloroplasts was detected in the range from 670 nm to 725 nm.

### 2.6 Promoter::GUS Transgenic Analysis

The promoter region from -677 to +11 of the genomic locus of AtDAT1 from Col-0 and Ler were amplified by PCR with the primer pair ProDAT1-SGW/ProDAT1-AGW (for sequences see Supplemental Table S1). The respective fragment was cloned into pENTR/D-TOPO and then into pMDC163 (Curtis and Grossniklaus, 2003), to be transformed into *Arabidopsis* by *Agrobacterium*-mediated gene transfer.

Histochemical staining of GUS activity was analyzed in plants of the T2-generation that had been germinated on liquid media. For GUS staining seedlings and adult plants were washed in sodium phosphate buffer and afterwards incubated overnight at 37°C in this buffer containing 1 mM X-Gluc (5-bromo-4-chloro-3-indolyl-beta-D-glucuronic acid) and 0.5 mM K_3_Fe(CN)_6_. Afterwards chlorophyll was removed for documentation by several washings with hot ethanol.

### 2.7 Recombinant Expression of AtDAT1 Variants in *E. coli*

*E. coli* strain BL21(DE3) RIL was transformed with cDNA of AtDAT1 variants in pGEX-2TM-GW (see above) and grown in LB medium with appropriate antibiotics until they reached an OD_600_ of 0.5. Then expression was induced by addition to a final concentration of 0.1 mM isopropyl-β-D-galactoside (IPTG) and the culture was grown for 20 h at 18°C. Afterwards cells were pelleted by centrifugation and washed once with TE buffer including 100 mM NaCl. After further centrifugation cells were resuspended in 20 mM Tris, pH 8, with Protease Inhibitor Cocktail from Biotool (Oberasbach, Germany). This suspension was sonicated and afterwards centrifuged with 18,000 x g to clear the crude extract from cell debris.

The recombinant His-tagged AtDAT1 protein variants from this crude extract were purified with Protino Ni-NTA agarose from Macherey-Nagel, (Weilmünster Germany) according to manufacturer’s protocol. Therefore, the column was equilibrated and loaded with 10 mM imidazole, washed with 20 mM imidazole, and elution of His-tagged proteins was achieved with 250 mM imidazole. Imidazole was removed by dialysis with Float-A-Lyzer Dialysis Device from Roth (Karlsruhe, Germany) in 10 mM potassium phosphate, pH 8. Protein content was determined with the Bio-Rad Protein Assay (Bio-Rad, München, Germany) according to manufacturer’s protocol. Specific detection of His tagged proteins on a western blot was achieved with a monoclonal His Tag antibody conjugated to alkaline phosphatase (antikoerper-online.de, Aachen, Germany).

### 2.8 Enzyme assays to determine D-AA specific aminotransferase activity

The standard reaction mixture with 2-OG as amino group acceptor contained D-Ala (10 mM), 2-OG (50 mM) and pyridoxalphosphate (PLP; 50 µM) in potassium phosphate buffer (100 mM, pH 8). For assays with pyruvate as amino group acceptor D-Ala and 2-OG were replaced by D-Met (10 mM) and pyruvate (50 mM), respectively. To determine substrate specificity, the tested D-AAs were all applied in 10 mM concentration. All assay reactions in triplicates were started by addition of 3-8 µg of purified protein, incubated at 37°C, and samples were taken at different time points up to 90 min. Each sample was derivatized and the amino acids measured as described below.

For the determination of K_M_ and V_max_ values different D-Met concentrations (0.1, 0.5, 1.0, 2.0, 5.0, 10.0, 20.0 and 50.0 mM D-Met) have been incubated with the enzyme AtDAT1 and pyruvate as cosubstrate (50 mM). Produced D-Alanine was analyzed after 0, 5 and 10 min. With the means of three biological replicates for any D-Met concentration and time point the slope of the time course was calculated and normalized to the protein amount used. To determine K_M_ and V_max_ values a linearization according to Hofstee (1959) was used.

### 2.9 Amino acid extraction and determination from plant material

Amino acid extraction and derivatization was performed as described before (Gördes et al., 2011). The incubation time of derivatization was elongated to 3 h and the derivatized liquid volume was adjusted with acetonitrile instead of methanol.

Almost all experiments were focused on the measurement of D/L-Alanine, D/L-Glutamate and D/L-Methionine. To determine and quantify these amino acids in plant extracts and enzyme assays standard materials were purchased from Sigma-Aldrich (Steinheim, Germany). Other chemicals were obtained in LC/MS grade from Roth (Karlsruhe, Germany). An Acquity–SynaptG2 UPLC-MS system from Waters (Manchester, England) was used for quantification, operated in positive electrospray ionization mode. The mass spectrometer was operated at a capillary voltage of 3000 V and a resolution of 20000. Separation of the amino acids was carried out on a Waters Acquity C_18_ HSS T3, 1.0 × 150 mm, 1.8 µm column with a flow rate of 50 µl/min and a 22 min gradient from 70 % water to 99 % methanol (both with 0.1 % formic acid). For quantification, 3 µl of sample were injected and a 5-point calibration from 0.125 µM to 1250 µM was used.

The quantification of malonyl-methionine ([M+H^+^] 218.022) and malonyl-ACC ([M+H^+^] 188.050) was performed relatively using the same LC/MS system described above. However, the stationary phase was changed into a Waters Acquity C_18_ HSS T3, 2.1 × 100 mm, 1.8 µm column, and a flow rate of 0.2 ml/min with a 15 min gradient from 99 % water to 99 % methanol (both with 0.1 % formic acid) was used for separation. The malonylated compounds were identified by the exact mass of their molecular ion.

### 2.10 Analysis of ethylene

For assaying ethylene production, *Arabidopsis* seedlings were grown in glass vials (18 ml) containing 3 ml solid medium (30 seedlings per vial) for six days. The vials were closed with rubber septa and opened once before measuring. After 30-90 min of further incubation ethylene accumulating in the free air space was measured by gas chromatography using a gas chromatograph equipped with a flame-ionization detector (Felix et al., 1991).

### 2.11 Statistical evaluation

Data was analyzed with IBM SPSS Statistics 24. Significances were analyzed using an independent two-sided Student’s t-test. For further analyses between and within genotypes, we used an ANOVA followed by Post hoc tests; Gabriel or Games-Howell, depending on the equality of variances. For testing the homogeneity of variances a levene test was applied.

## 3 Results

### 3.1 AtDAT1 as a candidate gene for D-AA metabolization

Initially, we observed the almost absence of both D- to L-AA and D-AA to D-Ala/D-Glu conversion in Ler in comparison to other ecotypes (Gördes et al., 2013). According to the transamination hypothesis, the mutation of at least one D-AA specific transaminase could be responsible for this metabolic phenotype. With AtDAAT1, one candidate protein had been previously identified biochemically as such an enzyme (Funakoshi et al., 2008). To investigate its role *in planta* we started to analyze T-DNA insertion lines of the corresponding gene (At5g57850; afterwards designated as *AtDAT1*) regarding their D-AA metabolization capability.

Homozygous plants for such insertion lines, SALK_011686 and SALK_111981 (denoted as *dat1-1* and *dat1-2*, respectively; Figure 1a), were isolated and propagated for further analyses (see Table S1 for primer sequences). RT-PCR analysis of *AtDAT1* expression displayed no transcripts in *dat1-1* and *dat1-2* mutants compared to the corresponding wild type (Col-0) (Figure 1b). However, the *AtDAT1* transcript level in Ler seedlings was similar to the wild type Col-0, what was also observed elsewhere (Lempe et al., 2005). As shown before (Gördes et al., 2011), feeding with D-Met caused the highest accumulation of D-Ala, D-Glu and its respective L-enantiomer in *Arabidopsis* seedlings. Therefore, mutant and corresponding wild type seedlings were grown for 14 days on liquid 1/2 MS medium in light, then supplemented with D-Met and subsequently analyzed for their AA contents. In sharp contrast to Col-0, both insertion mutants of *AtDAT1*, were neither able to produce D-Ala, D-Glu nor additional L-Met after application of D-Met. This AA profile was similar to that found in seedlings of the Ler accession (Figure 1c).

**Figure 1:**
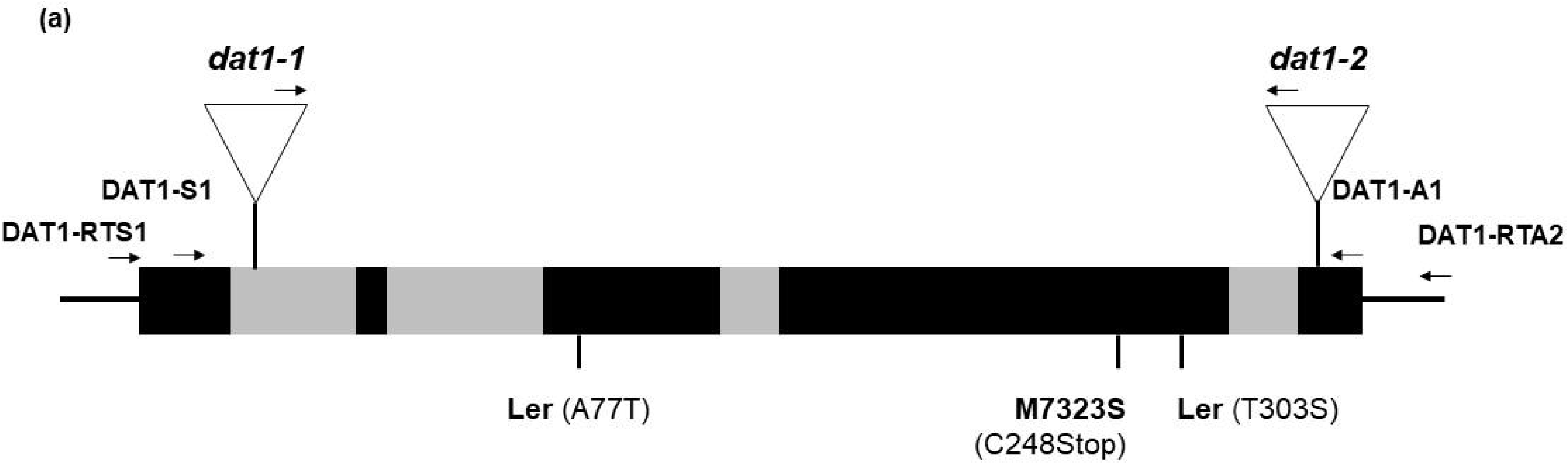
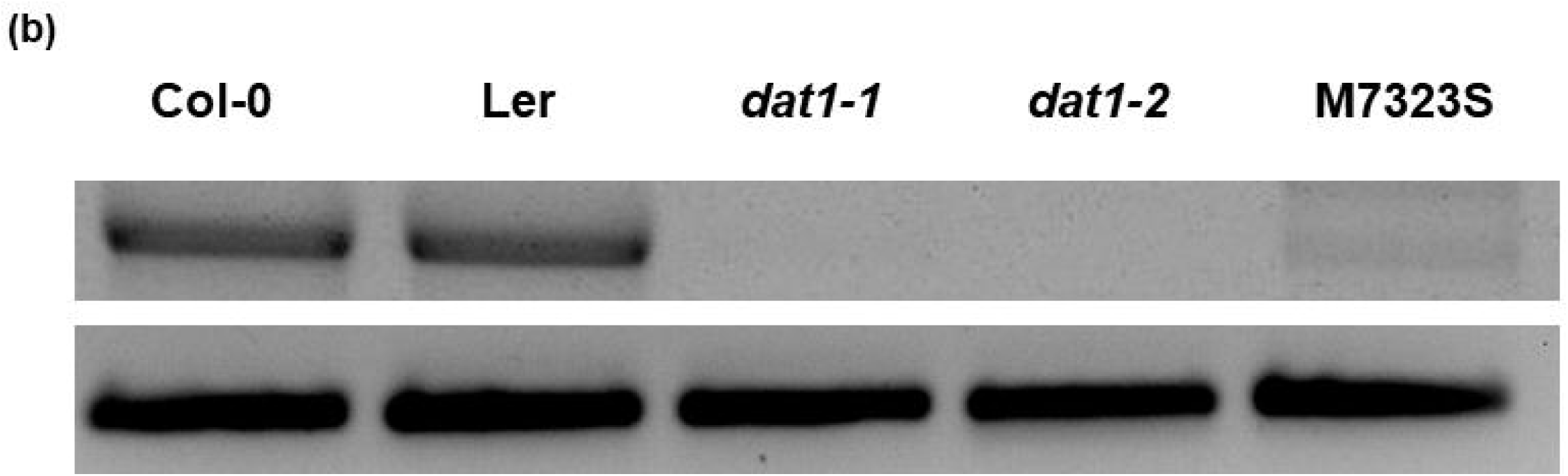
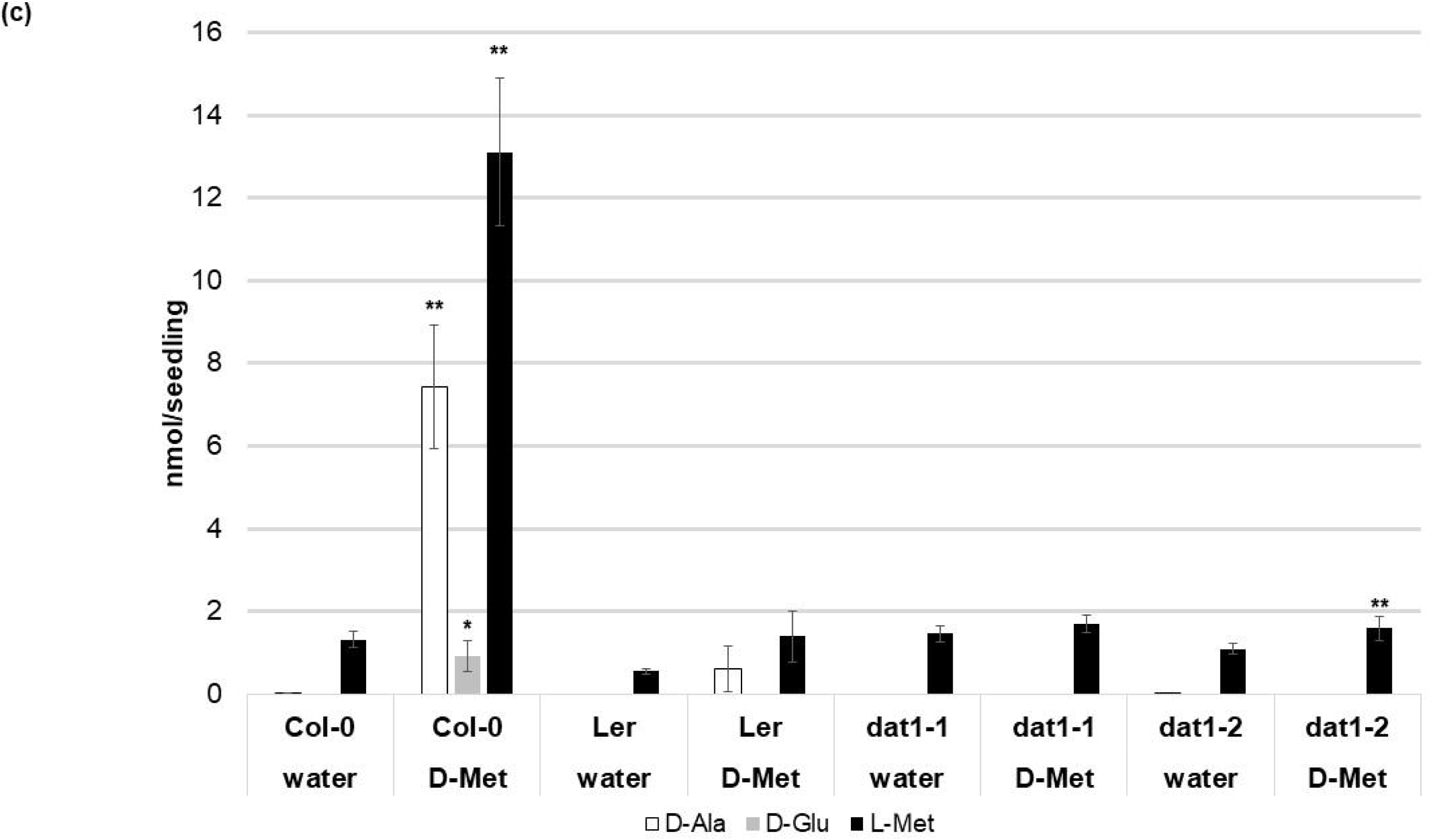
AtDAT1 as a candidate protein for D-AA metabolization in *Arabidopsis*. (a) Scheme of the genomic structure of AtDAT1 (exons and introns in black and grey, respectively) with the positions of T-DNA insertions in *dat1-1* and *dat1-2* as well as the mutations found in Ler and M7323S. Arrows indicate primers used for genotyping the T-DNA insertions and RT-PCR (for primer sequences see Table S1) (b) RT-PCR analysis of AtDAT1 expression in Col-0, Ler, *dat1-1, dat1-2* and M7323S (top: *AtDAT1*; bottom: *AtACT2*) (c) Contents of D-Ala (white), D-Glu (grey) and L-Met (black) in seedlings of Col-0, Ler, *dat1-1* and *dat1-2* without (water) and with D-Met treatment for 16 h (D-Met); For each measurement four seedlings were pooled and further processed. Error bars represent the standard deviation from three independent measurements. The asterisks indicate the significance level (t-test) of differences of all measurements to the respective line without D-Met treatment (* p<0.05; ** p<0.01).

Further *in silico* analyses of public transcriptomic data (Lempe et al., 2005) revealed that the accession M7323S displayed a strongly reduced *AtDAT1* transcript level, which could be confirmed by RT-PCR (Fig. 1b). When this accession was grown on D-Met supplemented medium defects in AA metabolism were observed (Figure S1) like in Ler and the *dat1* mutants. Sequencing of the genomic locus and the cDNA of *AtDAT1* from M7323S revealed that this gene contains a T→A mutation at genomic position +1259. This leads to a nonsense mutation at the third position of a cysteine codon (TGT) to a stop codon (TGA) at position 248 of the AA sequence (C248STOP) (Fig. 1a). In contrast, sequencing the genomic locus and the cDNA of *AtDAT1* from Ler just revealed two missense mutations leading to AA exchanges of the peptide sequence (A77T and T303S) (Fig. 1a).

To examine whether these mutations in the *AtDAT1* Ler allele are responsible for the metabolic aberrations in this accession, we performed different genetic approaches. Firstly, ubiquitin promoter-driven expression of the *AtDAT1* Col-0 allele in transgenic Ler plants led to the reconstitution of the D-Met metabolism in Ler and its complementation of the *dat1-2* mutant (Figure 2a). Secondly, F1 seedlings derived from crosses between Col-0 and Ler and between Col-0 and *dat1-2* displayed no D-Met metabolization defect as observed in Ler and *dat1-2*, irrespective of the maternal origin, whereas the offspring of Ler x *dat1-2* did (Fig. 2b). These data unequivocally prove the defect of AtDAT1 function in the Ler accession and *dat1-2* T-DNA insertion mutant.

**Figure 2:**
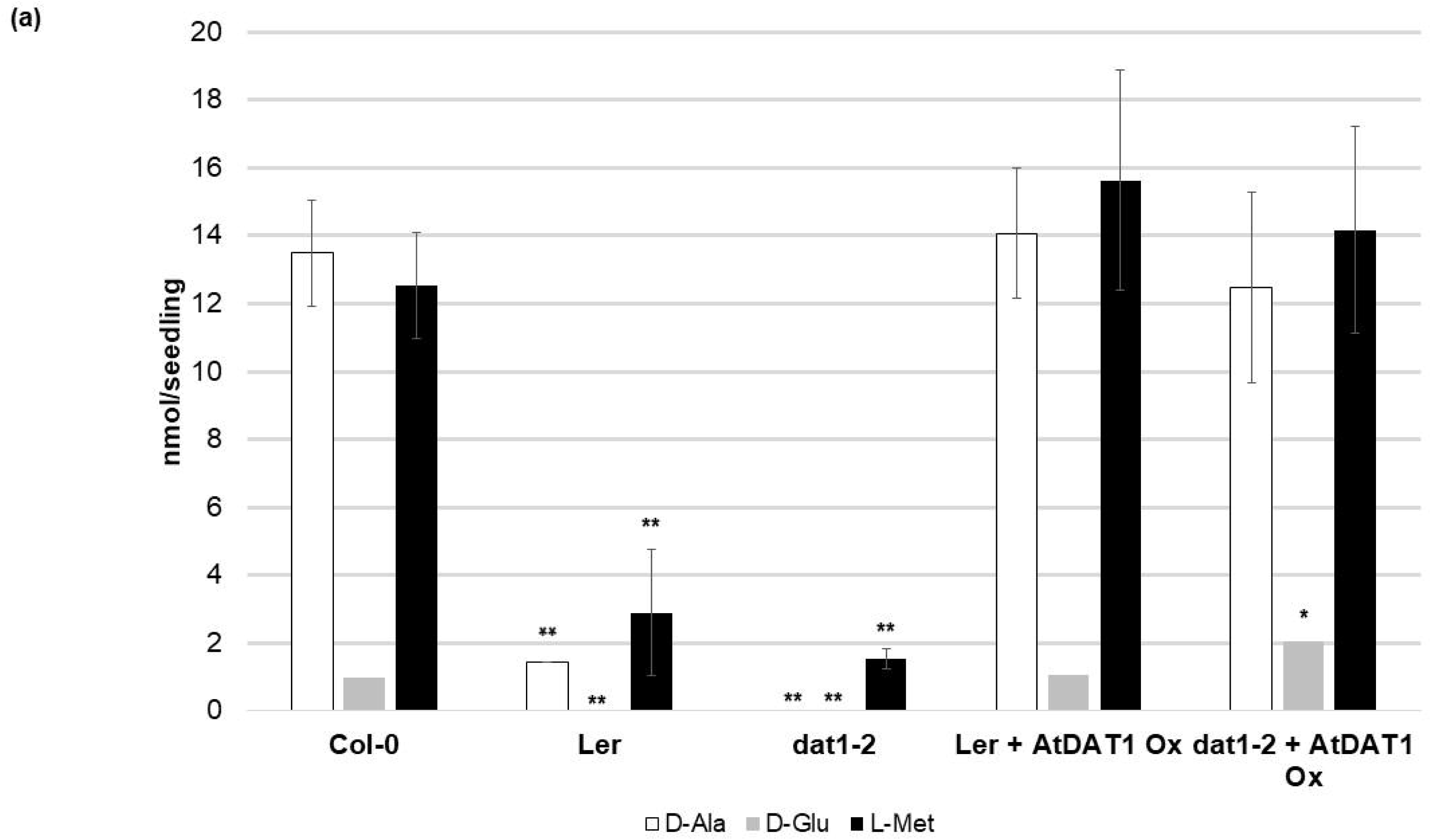
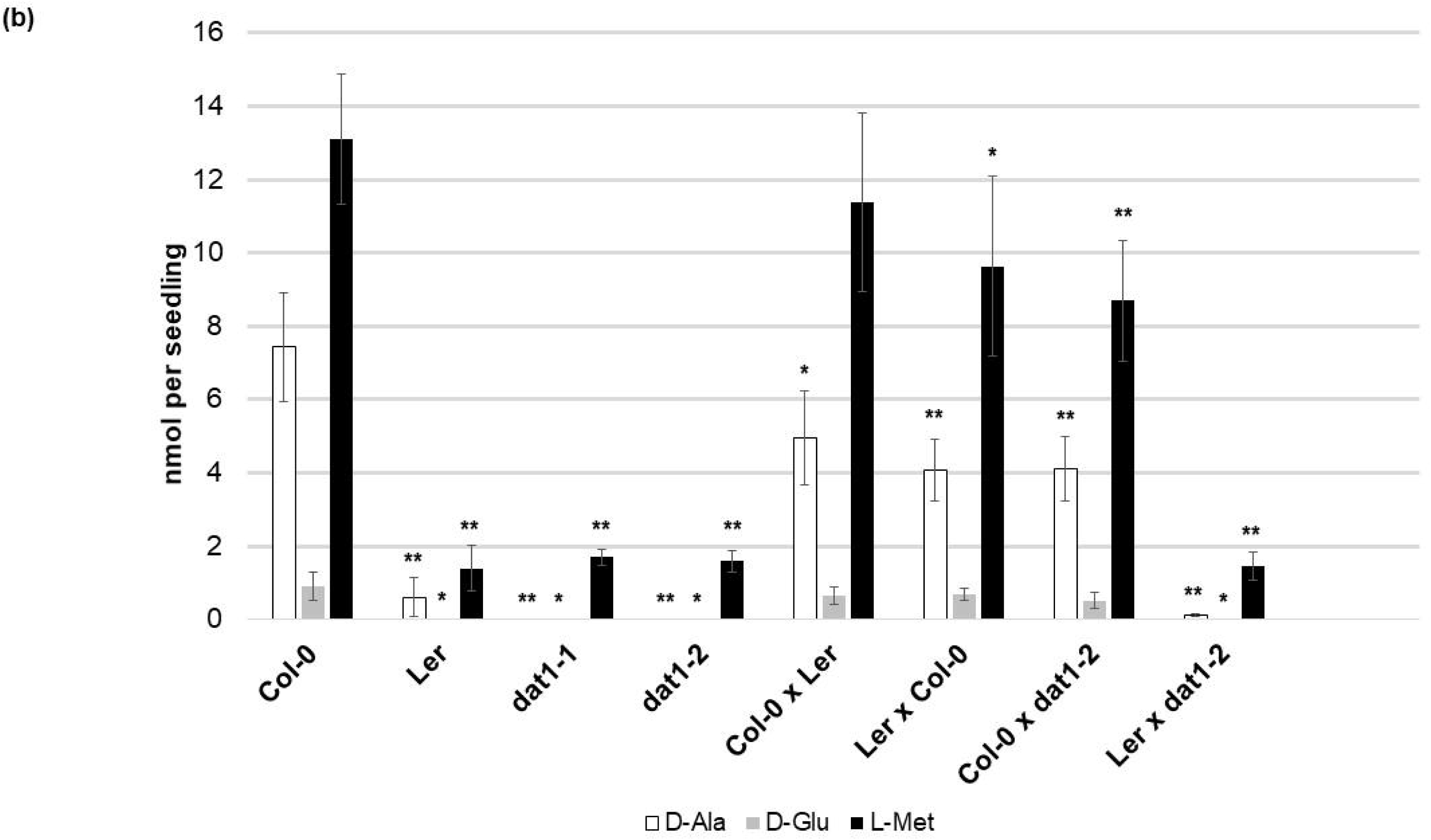
D-Met metabolization in lines overexpressing AtDAT1 and in F1 seedlings from crosses of Col-0, Ler and *dat1-2*. Contents of D-Ala, D-Glu and L-Met after overnight exposure to D-Met (a) in Ler and *dat1-2* overexpressing AtDAT1 (AtDAT1 Ox) and their corresponding background lines and (b) in F1 progeny seedlings of crosses of Col-0, Ler and *dat1-2* and their corresponding parental lines; for further information see Fig. 1c.

To answer the remaining question about the reason for this defect in Ler, the expression of AtDAT1 was analyzed. As mentioned before the transcript levels of this gene appeared similar in Col-0 and Ler (Figure 1b). This observation was supported by analysis of pDAT1::GUS plants (Figure S2a and S2b): There, it can be seen that this gene is expressed in seedlings and adult plants with less GUS staining in late floral stages and seeds (Figure S2a), corresponding to expression patterns displayed in the eFP browser (Winter et al., 2007). The activity of the *AtDAT1* promoters derived from Col-0 and Ler showed no apparent differences in seedlings irrespective of the presence of L-Met or D-Met in the media (Figure S2b). Subcellular mis-localization would have been another explanation for affected AtDAT1 function in Ler. Therefore, GFP-tagged AtDAT1 protein variants derived from cDNA of both ecotypes expressed under the control of the ubiquitin 10 promoter were transiently transformed into tobacco leaves (Figure S3). Both were found to be localized in all kinds of plastids. Therefore, a possible mis-expression or mis-localization caused by the mutation in the Ler allele of *AtDAT1* seems not to be the reason of its aberrant D-Met metabolization.

### 3.2 A missense mutation of the *AtDAT1* Ler allele leads to an almost complete loss of the enzymatic activity

To clarify if the enzyme encoded by the Ler *AtDAT1* allele is able to transaminate D-AAs, the Ler (AtDAT1_(Ler)_) and Col-0 (AtDAT1_(Col-0)_) versions of AtDAT1 were expressed with an N-terminal GST-tag in *E. coli*. After purification by affinity chromatography (for purification results see Figure S4) their enzymatic capabilities were tested according to Funakoshi et al. (2008).

To seek out the optimal substrate D-AA and experimental conditions, we initially tested AtDAT1_(Col-0)_ for its capability to transaminate 2-oxoglutarate (2-OG), like in Funakoshi et al. (2008), or pyruvate using 16 different D-AAs as amino group donors. With 2-OG used as amino group acceptor, a transaminase reaction was only detectable for the donors D-Met, D-Trp and D-Ala (Table S2), whereas with pyruvate as acceptor almost all applied D-AAs (with the exception of D-Pro) led to the formation of D-Ala (Figure 3). Furthermore, we measured an over 100 times higher activity for the enzymatic reaction with pyruvate as acceptor than with 2-OG, irrespective of the D-AA used as amino group donor (Table S2). The comparison of the AtDAT1_(Col-0)_ activities using different D-AAs and pyruvate as substrates revealed that D-Met was the best tested amino group donor (Figure 3). Using these two compounds as substrates we determined the K_M_ and V_max_ of AtDAT1_(Col-0)_ to be 17.4 mM and 0.07 nkat, respectively.

**Figure 3:**
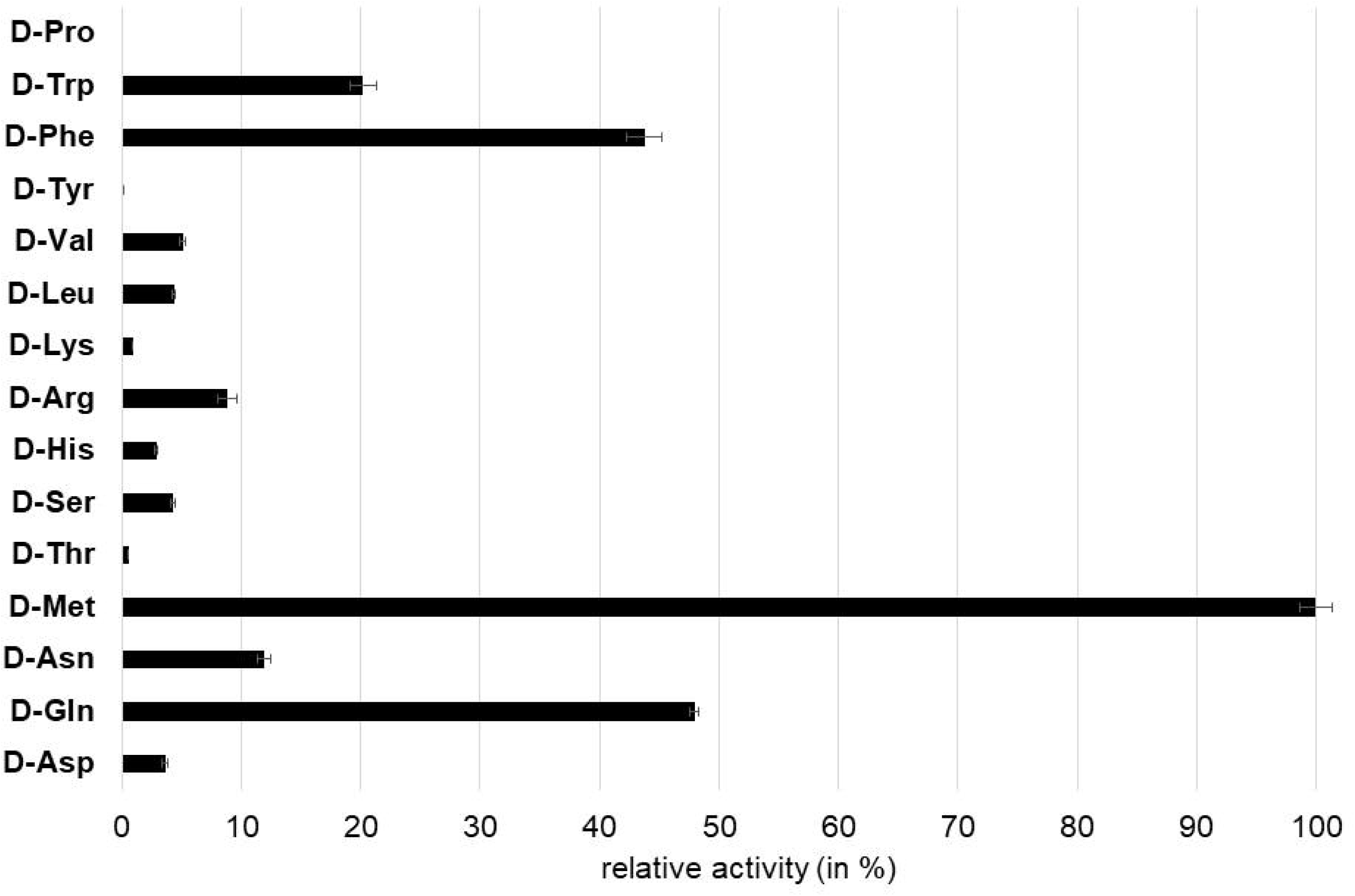
Relative D-Ala producing activity of AtDAT1 with different D-AAs as amino group donor and pyruvate as acceptor. Activity of reaction with D-Met was set to 100% and all other reactions were calculated in relation to it. Each bar represents the mean of measurement of three independent assays. Error bars (± SD).

To investigate the enzymatic activity of AtDAT1_(Ler)_ assays were performed with two substrate combinations: Firstly, with D-Met and pyruvate as amino group donor and acceptor, respectively, as the best substrate combination for AtDAT1_(Col-0)_ and, secondly, with D-Ala and 2-OG to evaluate previously published data (Funakoshi et al., 2008). As shown in Figures 4a and 4b for both substrate combinations the activity of AtDAT1_(Ler)_ dropped to 0-5% compared to AtDAT1_(Col-0)_.

**Figure 4:**
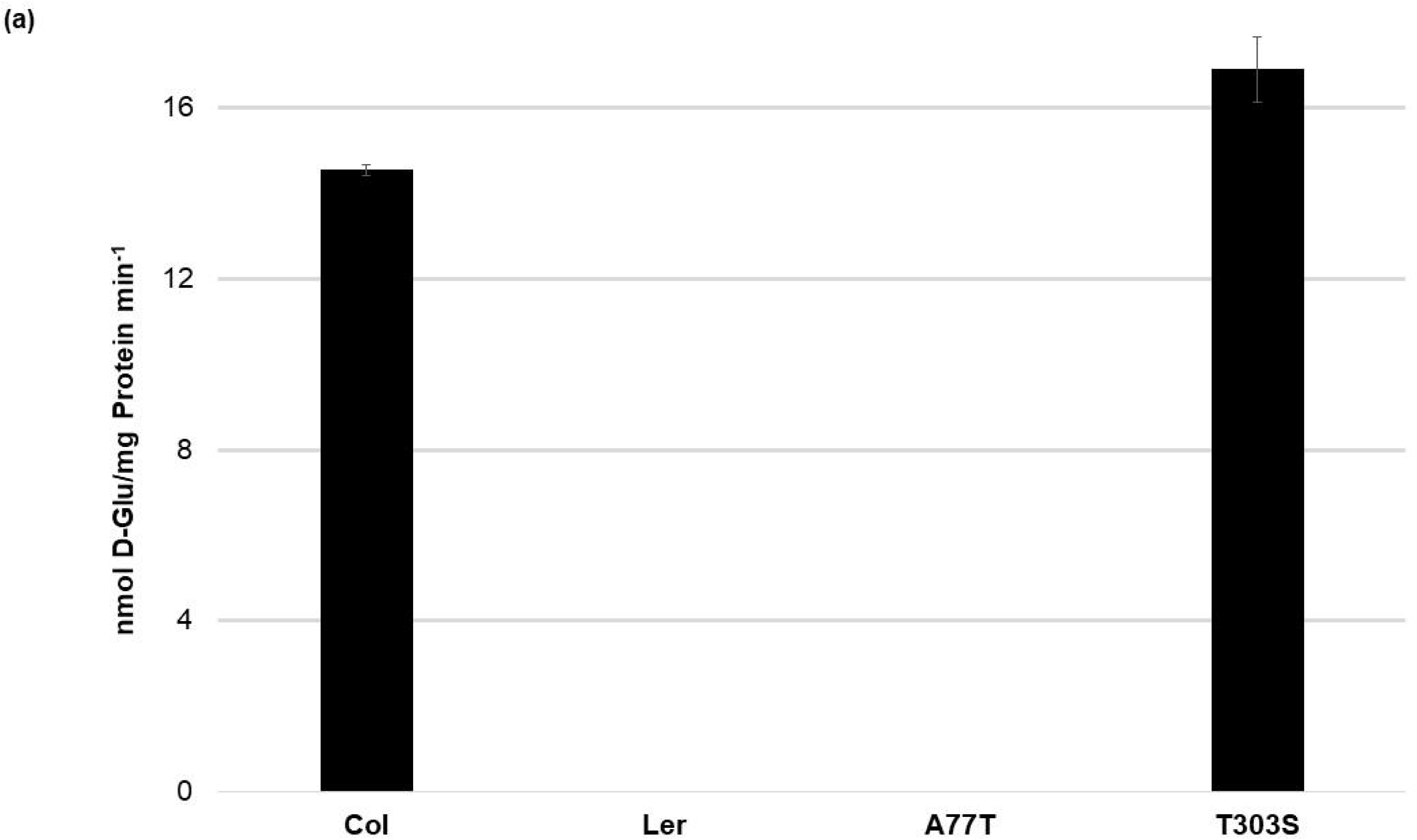
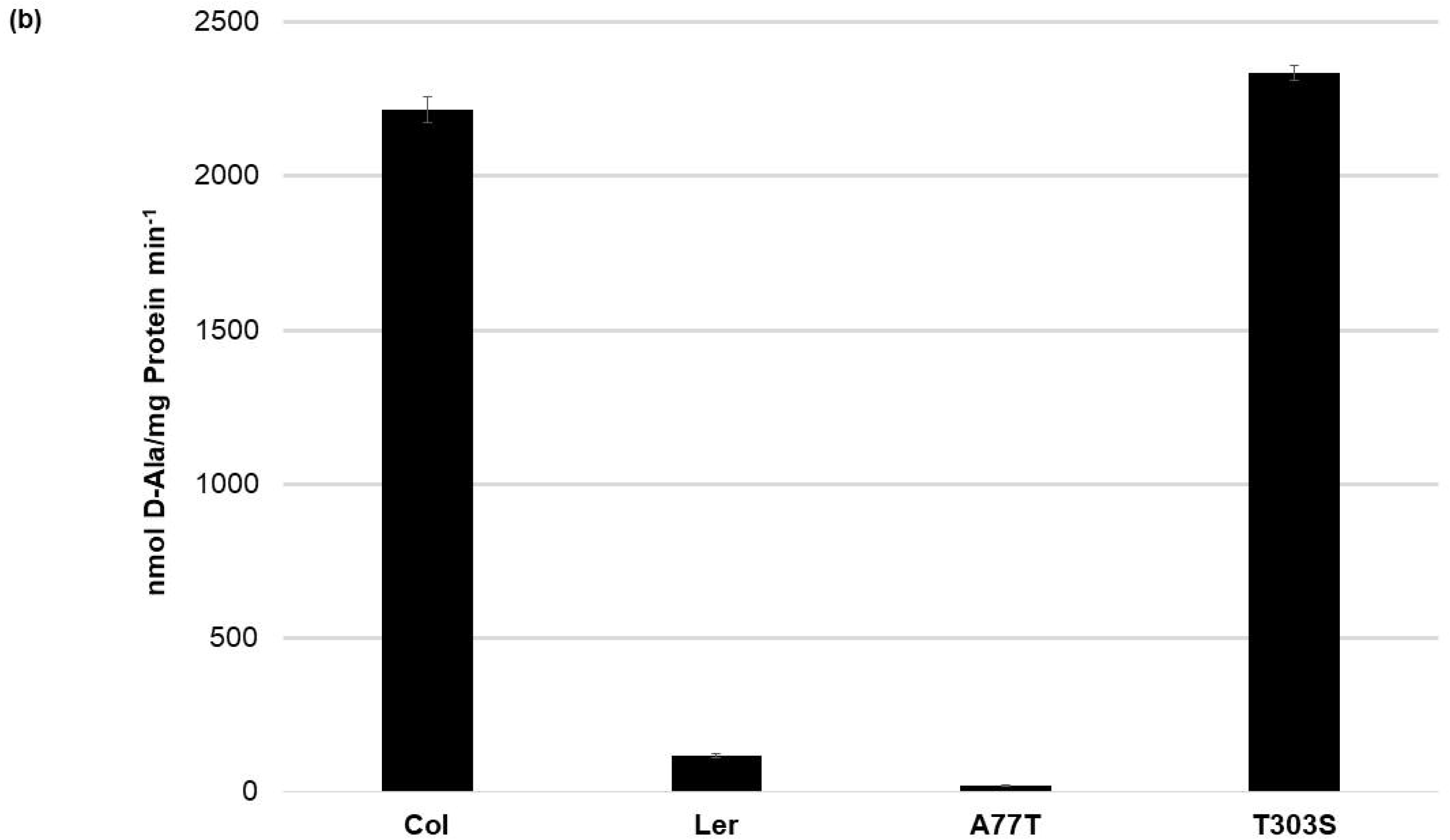
Activities of AtDAT1 variants. Transaminase activities of AtDAT1_(Col-0)_, AtDAT1_(Ler)_, AtDAT1_(A77T)_, and AtDAT1_(T303S)_ with D-Met as amino group donor and (a) 2-oxoglutarate or (b) pyruvate as acceptor molecule are displayed; for further information see Fig. 3.

At that point the question arose if just a single missense mutation (A77T and T303S) was sufficient to cause this activity loss in AtDAT1_(Ler)_. The alignment of DAT1 amino acid sequences from different plant species revealed that the alanine at position 77 is more conserved than the threonine at position 303 (Figure S5). To analyze the impact of the mutations of AtDAT1_(Ler)_, AtDAT1_(Col-0)_ derived isoforms harboring single amino acid exchanges were also expressed as N-terminal GST fusions in *E. coli*, affinity-purified and tested for their activity. Whereas the enzyme isoform with the T303S amino acid exchange AtDAT1_(T303S)_ showed an activity comparable to AtDAT1_(Col-0)_ (Figures 4a and 4b), the mutation A77T led to a strong decrease in the production of D-Glu (Figure 4a) and D-Ala (Figure 4b) with 2-oxoglutarate or pyruvate as substrates, respectively. Instead, the enzymatic defect of AtDAT1_(A77T)_ was quantitatively similar to that of AtDAT1_(Ler)_. From these data we concluded that solely this amino acid exchange is responsible for the activity loss of AtDAT1 in Ler. But the data also revealed that the Ler variant is not completely inactive with about 5% remaining activity in comparison to Col-0 (Figure 4b).

### 3.3 The loss of AtDAT1 leads to decreased seedling growth in response to D-Met

After identification of AtDAT1 as a central enzyme for metabolization of D-AAs, the question arose, to which physiological effects in *Arabidopsis* growth and development the loss of *AtDAT1* gene function would lead. Under greenhouse conditions in soil growth of *dat1-1* and *dat1-2* mutant plants could not be distinguished from Col-0 (Figure S6).

An obvious question in this respect was how these mutant lines would grow in presence of D-Met. Germination on D-Met in concentrations of 500 µM affected *dat1-1* and *dat1-2* seedling growth significantly compared to the corresponding wild type, whereas Ler took an intermediate position (Figure 5a). Testing this growth behavior in the dark revealed an even more pronounced growth difference between *dat1* mutants and Col-0 (Figure 5a). All these growth differences were specific for D-Met, whereas the addition of the same concentrations of L-Met did not lead to these differential effects (Figure 5a). Altogether, D-Met inhibited seedling growth specifically in AtDAT1 affected lines.

**Figure 5:**
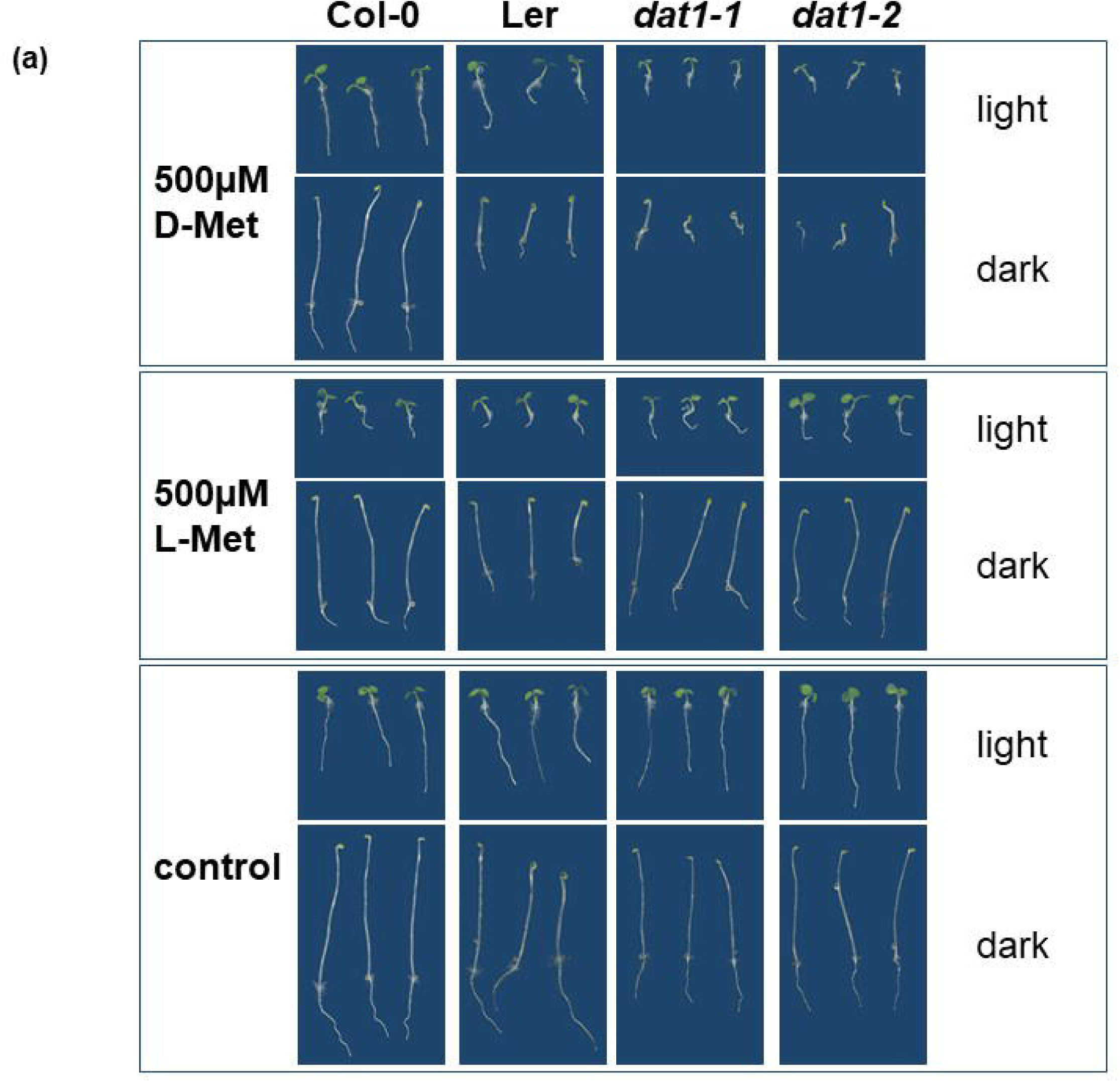
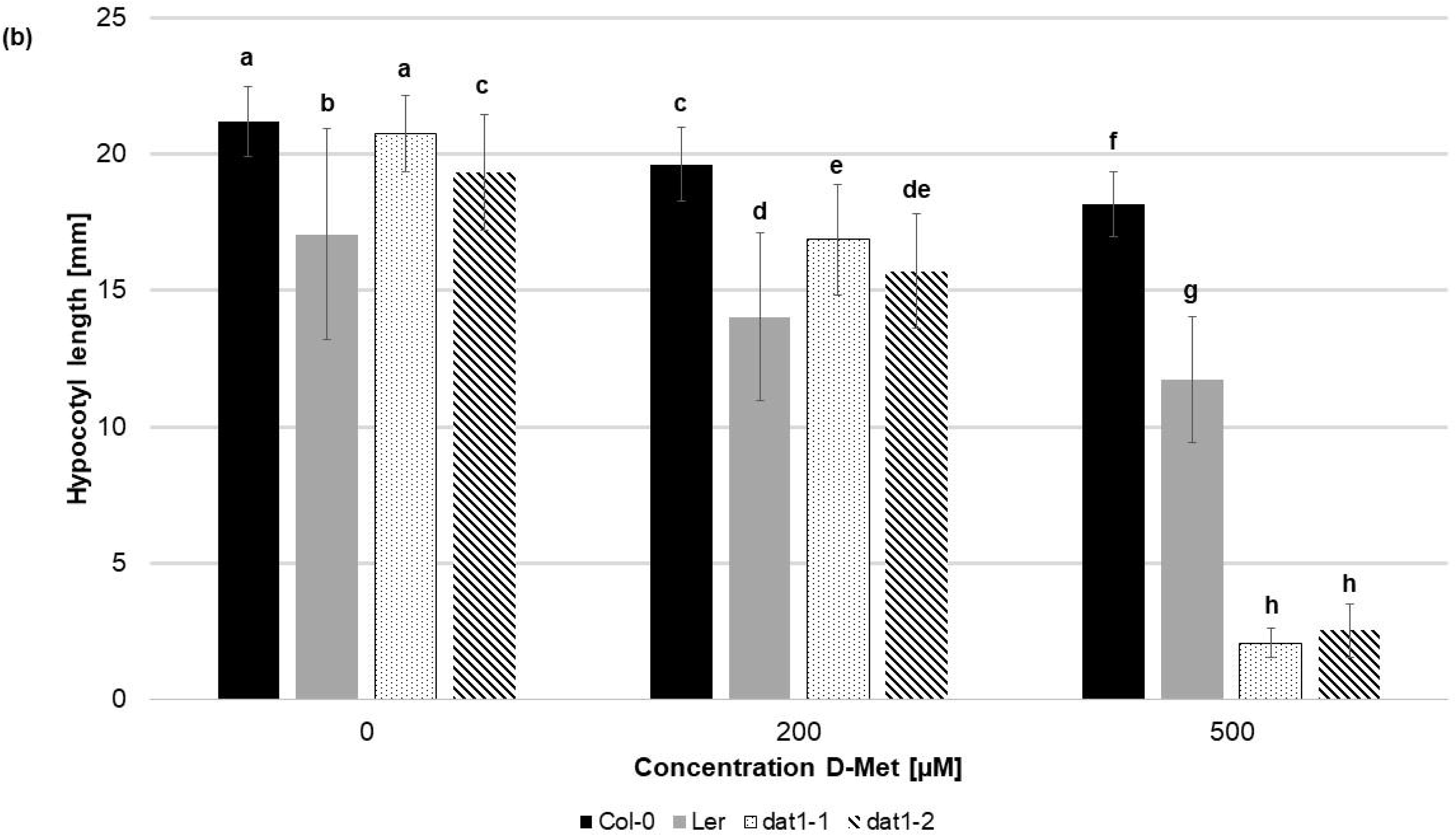
Seedling growth is differentially suppressed by D-Met in *AtDAT1* knock out-lines. (a) Seeds of Col-0, *dat1-1*, and *dat1-2*, and Ler were germinated either in permanent light (light) or in darkness (dark) on different solid growth media (without supplementation, with 500µM D-Met or with 500µM L-Met supplemented). Afterwards, hypocotyl growth of the dark grown plants was measured (b). Bars (Col-0: black, Ler: grey, *dat1-1*: dotted, *dat1-2*: striped; n=30) represent the average hypocotyl length. Different letters indicate statistically significant differences (p<0.05) tested by an ANOVA. Error bars (± SD).

### 3.4 *Atdat1* mutants display enhanced D-AA stimulated ethylene production

Especially the growth effects of *dat1-1* and *dat1-2* in the dark (Figure 5a) indicate a triple response caused by the gaseous plant hormone ethylene. This gets even clearer with a look on the hypocotyl length of the four dark grown lines (Figure 5b): There was a highly significant decrease of *dat1-1* and *dat1-2* hypocotyl length of about one eighth compared to Col-0 grown on 500 µM D-Met. Increasing L-Met concentrations also led to shorter hypocotyls, but to similar extent in mutants and wild type plants. Furthermore, the growth inhibition was by far not as strong as with D-Met (Figure S7).

To elucidate if this observation was really caused by ethylene we measured its production in Ler, *dat1* mutants and Col-0 grown in the dark. As it can be seen in Figure 6a the addition of 500 µM D-Met was sufficient to induce a significant increase of up to threefold of ethylene production in light grown Ler and *dat1* mutants compared to Col-0. Even stronger changes in ethylene production could be observed for both *dat1* mutant lines germinated in the presence of D-Met also in the dark, whereas Ler displayed again an intermediate reaction (Figure 6b).

**Figure 6:**
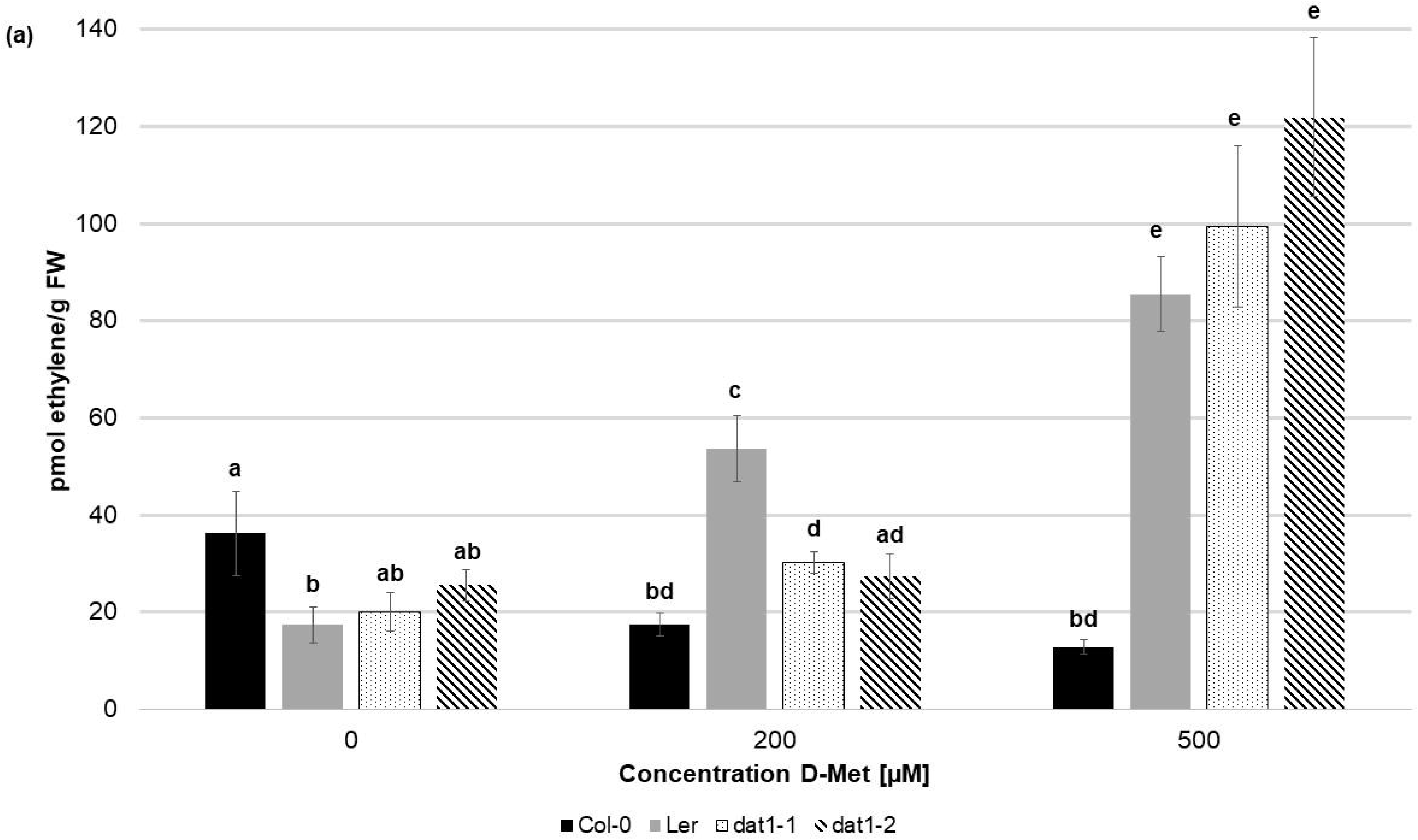
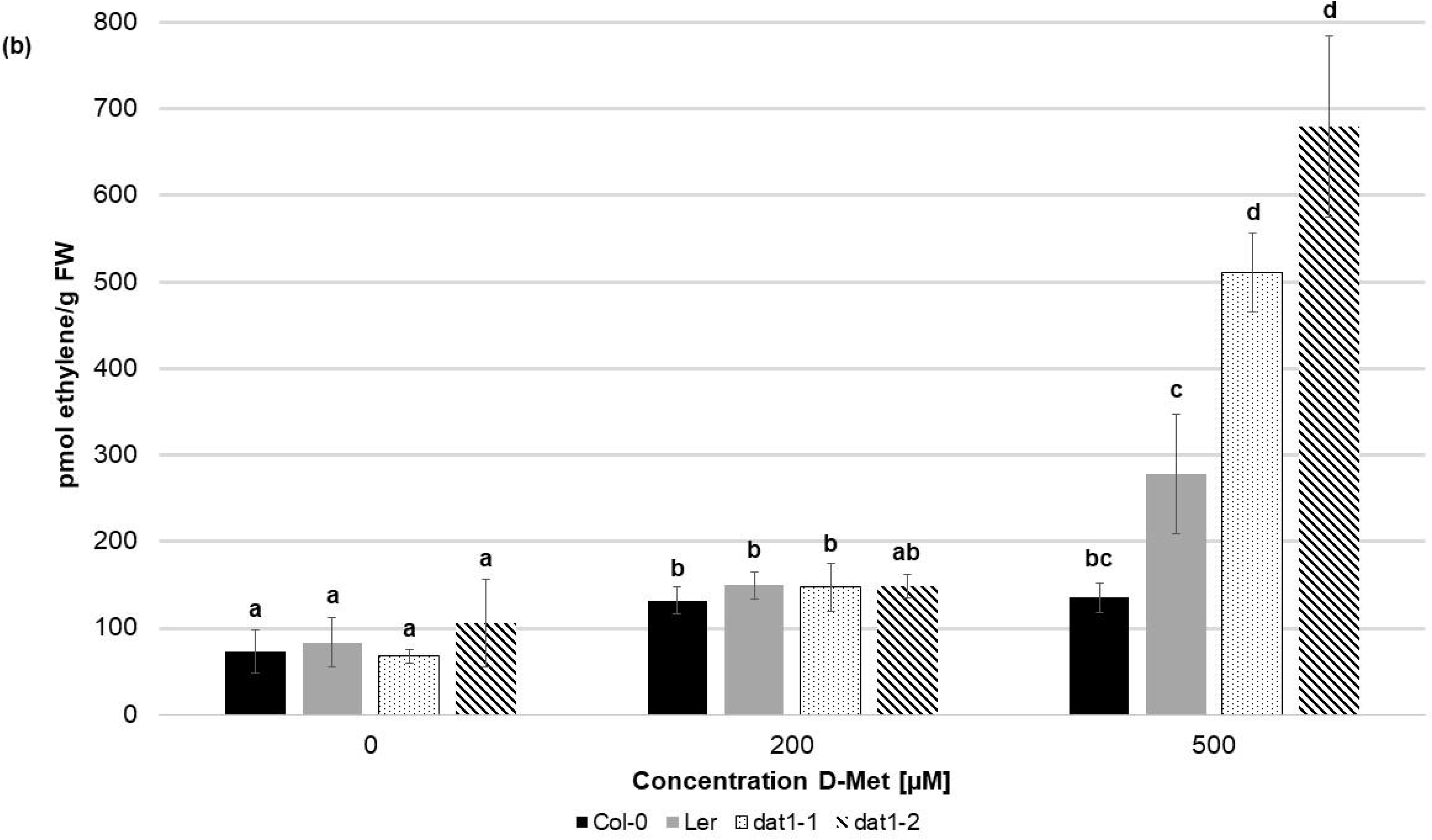
D-Met leads to increase of ethylene in AtDAT1 knock out-lines. Ethylene contents in seedlings of Col-0, Ler, *dat1-1*, and *dat1-2* were measured after growth (a) in permanent light or (b) in permanent darkness in vials with solid growth media supplemented with 200µM and 500µM D-Met, and additionally without supplementation (control). The bars represent averages of three biological replicates; for further information, see Fig. 5b.

As mentioned above the increase of ethylene production by D-AAs was attributed to competitive malonylation of D-AAs instead of ACC, which should lead to enhanced ACC oxidation resulting in higher ethylene concentration (S F Yang and Hoffman, 1984). To verify this assumption, we measured the contents of malonyl-methionine and malonyl-ACC in D-Met treated seedlings. In these measurements we detected a significant increase of malonyl-methionine in Col-0, Ler and *dat1* seedlings upon D-Met treatment (Figure 7a and 7b). This accumulation was far higher (up to 5 fold) in both mutants compared to the corresponding wild type, irrespective of the light regime (Figure 7a and 7b). Furthermore, Ler showed also a D-Met induced overaccumulation of malonyl-methionine in the light (Figure 7a), but not in darkness (Figure 7b).

**Figure 7:**
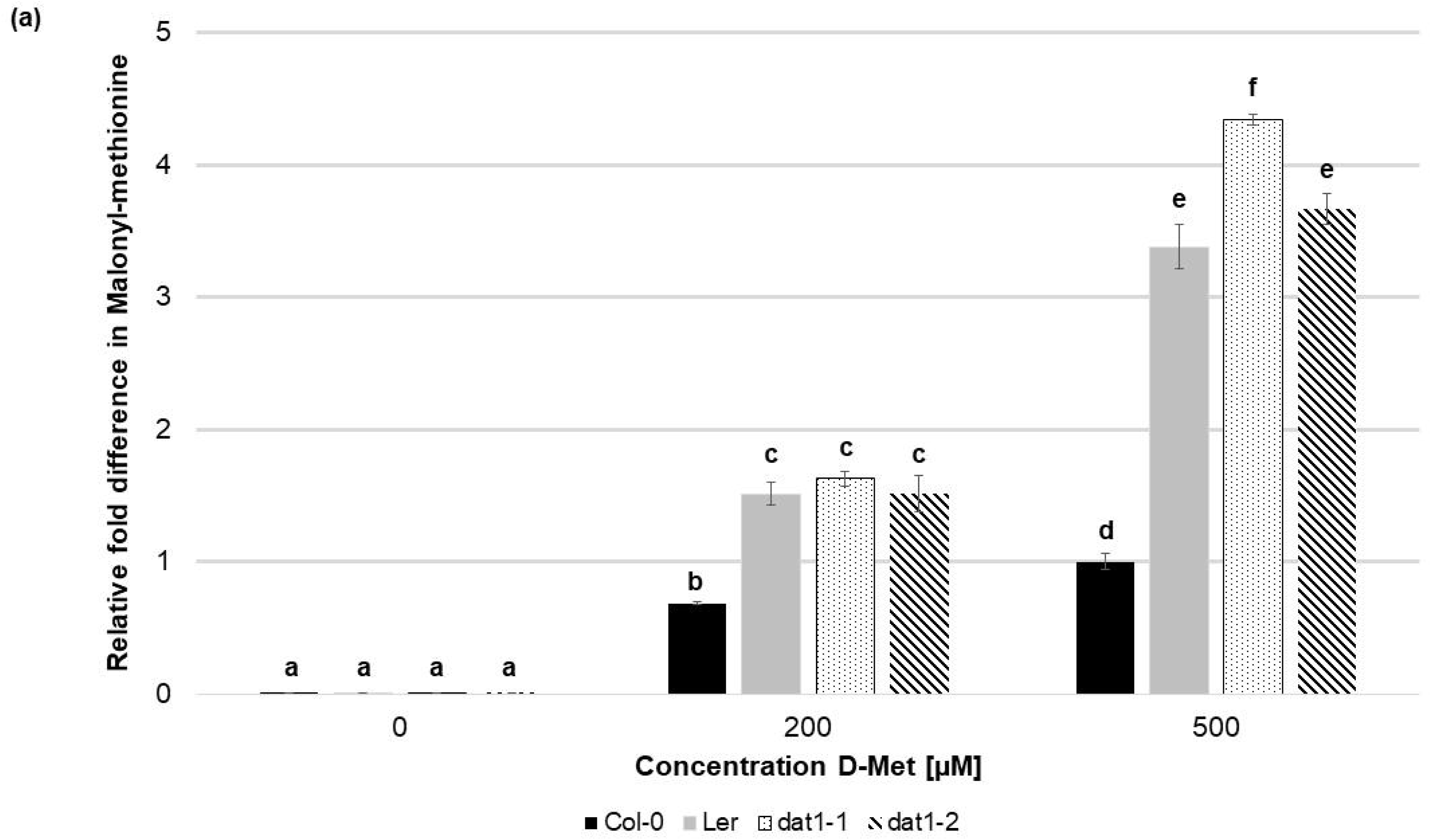
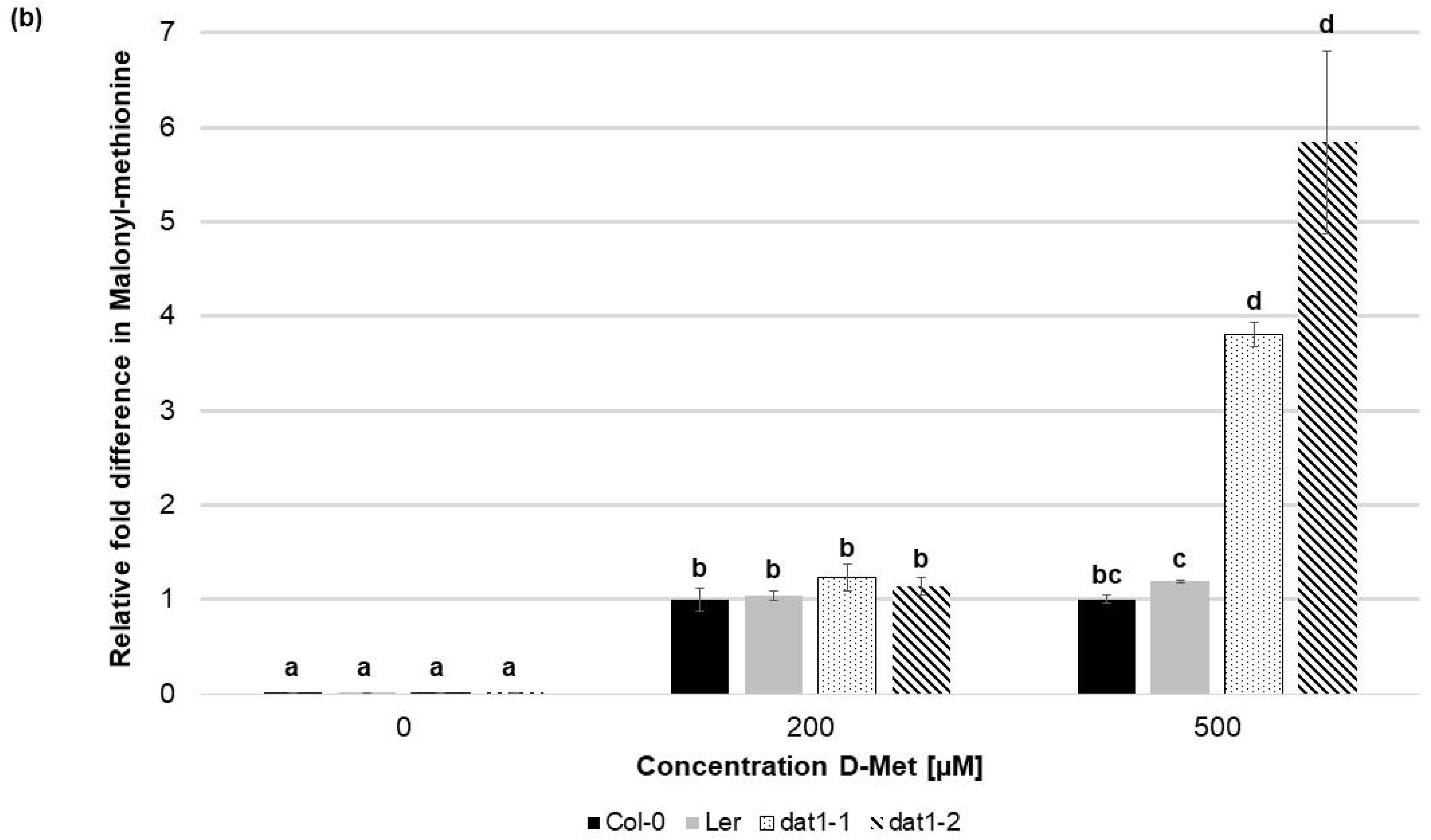
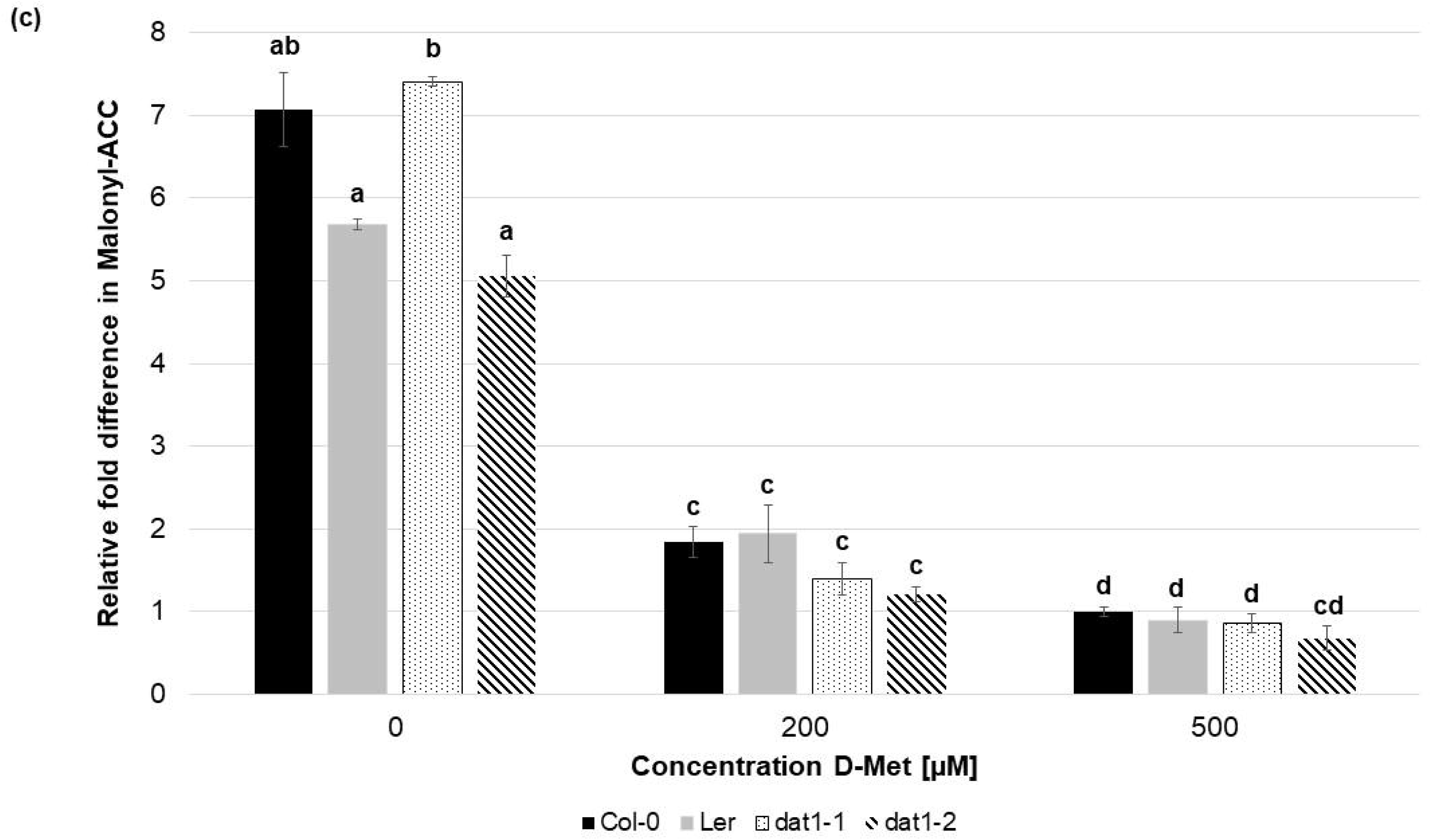
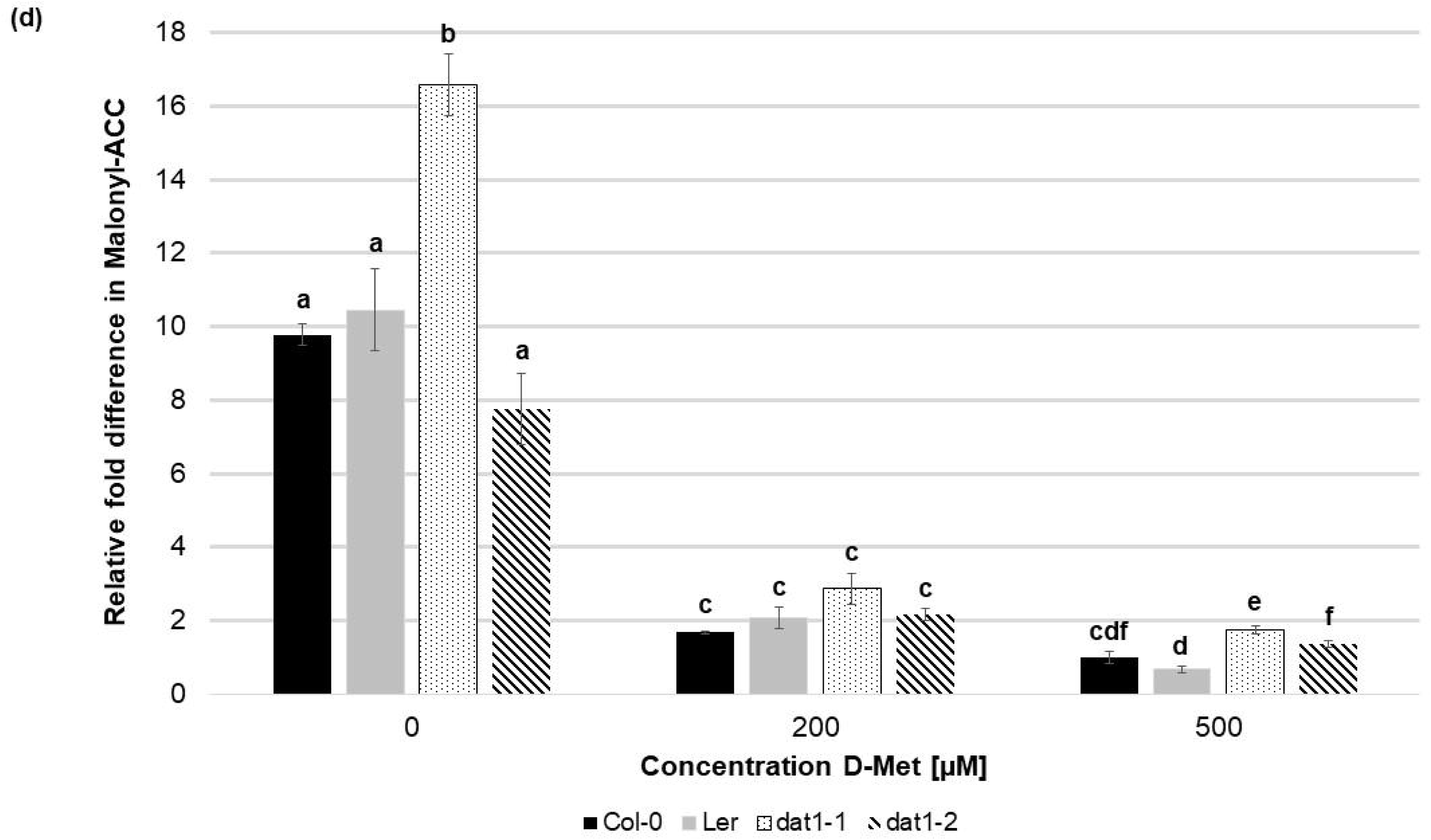
D-Met affects formation of malonyl-methionine and malonyl-ACC differently in Col-0, Ler and *dat1* mutants. Malonyl-methionine contents in seedlings of Col-0, Ler, *dat1-1*, and *dat1-2* were measured after growth (a) in permanent light or (b) in permanent darkness grown on agar plates supplemented with 200µM and 500µM D-Met, and additionally without supplementation (control). The relative values are given in fold changes with the values of Col-0 at 500µM D-Met set to 1. Furthermore, the relative malonyl-ACC contents in these seedlings were measured after growth on the given D-Met concentrations plus 10 µM ACC. Seedlings were either grown (c) in permanent light or (d) darkness; for further information, see Fig. 5b.

Since the amount of malonyl-ACC in these experiments was below our detection limit, we added 10 µM ACC to the media and measured once more malonyl-ACC in all lines. In this case we were able to detect large amounts of malonyl-ACC in all treated lines, which decreased drastically upon D-Met addition (Figure 7c and 7d). D-Met induced malonyl-ACC reduction was not due to production of malonyl-methionine caused by ACC, which was comparable with and without ACC addition (Table S3). Nevertheless, there was no significant difference of malonyl-ACC reduction of Ler and *dat1* mutants to Col-0 at higher D-Met concentrations (Figure 7c and 7d).

## 4 Discussion

For several decades the detrimental, but partially also beneficial, effects of D-AAs on plants have been investigated (Valdovinos and Muir, 1965;Aldag and Young, 1970;Erikson et al., 2004;Erikson et al., 2005;Gördes et al., 2011;Hill et al., 2011). It is noteworthy, that there are reports of some D-AAs synthesized *de novo* by plants (Brückner and Westhauser, 2003;Strauch et al., 2015). Especially in recent years there was growing evidence that almost all D-enantiomers of proteinogenic L-AAs are taken up by plants (Aldag and Young, 1970;Forsum et al., 2008;Gördes et al., 2011;Hill et al., 2011), and also metabolized in significant amounts (Aldag and Young, 1970;Gördes et al., 2011). With the proof provided, the long standing question was addressed how D-AAs are metabolized in plants.

In the light of the observations of Gördes et al. (2011), three possible mechanisms for this process had been suggested: racemization, deamination and transamination of D-AAs (Vranova et al., 2012;Gördes et al., 2013). Our data indicate that transamination by AtDAT1 is responsible for major steps of D-AA turnover in *Arabidopsis*, which is reflected by its broad range of D-AA specificity (Figure 3). Furthermore, we showed that the major product of this enzymatic reaction is D-Ala with D-Met as the favored amino group donor as it was also the case when plants were fed with D-AAs. Why D-Ala was known as a primary product of D-AA metabolization in plants could also be answered by our studies. This effect is caused by the property of AtDAT1 to prefer pyruvate over 2-OG (Table S2). In comparison to the work of Funakoshi et al. (2008), who used 2-OG as amino group acceptor for their characterization of AtDAT1, our results revealed a higher V_max_ with pyruvate as substrate. Most interestingly, the different enzymatic activities with pyruvate and 2-OG as amino acceptors with ratios of 100:1 and more (Table S2) were in a comparable range as the D-Ala/D-Glu ratios found in plants after D-AA application (Gördes et al., 2011). This preferential accumulation of D-Ala in D-AA treated plants is in accordance to the presented preference for pyruvate as substrate of AtDAT1.

A major question in our studies was the role of AtDAT1 in D-AA stimulated ethylene production. As it could be shown in this report this phenomenon is tightly connected to AtDAT1. The loss of this protein leads to a significant increase of ethylene after D-Met application (Figure 6), resulting primarily in shortening of the hypocotyl of *dat1* mutants irrespective of the light regime (Figure 5). This treatment led also to an increased production of malonyl-methionine, especially in the *dat1* mutants, and the content of malonyl-ACC developed reciprocally in all tested lines (Figure 7). Especially, the reciprocal malonylation capacity of D-Met and ACC implies that this is the reason for the observed increase in ethylene production and enhanced triple response. But this conclusion has to be reviewed critically, because no differences in ACC malonylation could be detected between D-Met treated Col-0 and *dat1* mutants. At this point it must be noted, that in all previous studies the production of ethylene in response to malonylation of ACC and D-AAs had been measured in overnight feeding experiments. But the physiological growth responses and the contents of the plants over a longer period of the treatment, like in the present study, have been shown in this study for the first time.

The increased production of malonyl-methionine and ethylene without a decreased malonyl-ACC production in *dat1* mutants in comparison to the control raises the question if the original working model of D-Met stimulated ethylene production needs an additional factor to explain our observations. Such an explanation may be that malonylation is not the only way to regulate the ACC level in plants. They are also able to conjugate ACC with glutathione to γ-glutamyl-ACC (GACC) and with jasmonic acid to JA-ACC to control the ACC content. Additionally, plants can degrade ACC irreversibly by an ACC deaminase α-ketobutyrate (for reviews about ACC content regulation see Van de Poel and Van Der Straeten (2014), Le Deunff and Lecourt (2016), and Vanderstraeten and Van Der Straeten (2017)). To date neither the contribution of each of these ACC degradation pathways to control the ACC pool nor their interplay has been studied, yet. It remains to be investigated if D-Met, its malonylated form or the loss of this pathway have an impact also on the alternative ACC degradation pathways.

Another explanation for the contradicting results may be given by the products of enzymatic activity of AtDAT1 on other enzymes. As it has been demonstrated in this study the loss of this enzyme leaves *dat1* mutants without the ability to produce D-Ala, D-Glu and additional L-Met in response to D-Met. Most interestingly, it was shown previously that D-Ala inhibits the ACC oxidase (Gibson et al., 1998;Brunhuber et al., 2000;Charng et al., 2001;Thrower et al., 2006) which catalyzes the last step from ACC to ethylene. This means that plants with functional DAT1 would malonylate D-Met instead of ACC. But the produced D-Ala would partially inhibit ACC oxidase and as one consequence the additional ACC would just be partially converted to ethylene. In *dat1* mutants this limiting effect of D-Ala would be lost and may explain the ethylene increase in these lines in comparison to the corresponding wild type. The same lack of D-Ala production of *dat1* mutants may also contribute to the higher content of malonyl-methionine in the mutants, but also to the comparable amount of malonylated ACC in all tested lines: It had been shown before that D-Ala also partially inhibits the putative malonyl transferase (Kionka and Amrhein, 1984;Liu et al., 1985;Chick and Leung, 1997). The malonylation of D-Met would then be limited by D-Ala in Col-0 but not in DAT1 affected lines. If D-Met is the preferred substrate of the malonyl transferase, the lack of significant differences between the tested lines in ACC malonylation would not be surprising. To confirm these assumptions that D-Ala influences the D-Met stimulated ethylene production by inhibiting the ACC oxidase, the ACC malonyl transferase or in combination of both, further physiological experiments with D-Ala and structural analogs like D-cycloserine are needed. But final answers to this question would be awaited by the identification of the ACC malonyl transferase and its biochemical and physiological characterization.

Undoubtedly, AtDAT1 is a key enzyme of D-Met stimulated ethylene production and seems to have quite specific effects in this regard. But the question remains where this D-Met comes from because it has never been reported in plants until to date. In contrast, it has been shown previously that D-Met is released by bacterial biofilms (Kolodkin-Gal et al., 2010;Vlamakis et al., 2013) and that different rhizosphere colonizing bacterial species are able to utilize D-Met as sole carbon and nitrogen source (Radkov et al., 2016). Biofilm formation on root surfaces as a bacterial pathogen protection strategy has been reported before (for a review see Vlamakis et al. (2013)). Possibly, AtDAT1 is part of bacterial biofilm recognition and therefore may be involved in plant-bacterial interaction.

This possibility would also offer an explanation why AtDAT1 is dispensable in particular *Arabidopsis* accessions as it has been shown for Ler and M7323S in this report. In a habitat without D-Met releasing bacteria in the rhizosphere, a recognition system for this compound would be also dispensable for the plant. But this relation needs to be proven first. Interestingly, DAT1 encoding genes seem to be found in almost all sequenced plant genomes (for a selection see Figure S5), and ethylene production in other plant species than *Arabidopsis* is also induced by other D-AAs like D-Leu, D-Thr, D-Val or D-Phe (Satoh and Esashi, 1980;1982;Liu et al., 1983). On the one hand it would be interesting if also other D-AAs than D-Met cause differential growth effects and ethylene production in *dat1* mutants. On the other hand, it should be tested if DAT1 proteins from different species have altered substrate specificities and therefore contribute to the adaptation of plants to changing microbial environments.

## 6 Author Contributions

ÜK and JS designed the study. JS and CH conducted most of the experiments and contributed equally to the study. VAL and SH conducted another part of the experiments. CH and MS analyzed most of the biochemical data and ÜK wrote the manuscript.

## 8 Acknowledgments

We would like to acknowledge Prof. Klaus Harter (Center for Plant Molecular Biology, University of Tübingen, Germany) for the provision of research facilities and critically reading the manuscript. Furthermore, we would like to thank to Nina Glöckner and Friederike Wanke for their excellent support in confocal fluorescence microscopy and to Georg Felix for his support in ethylene measurements.

