## Supplementary figures and images for "AtDAT1 is a key enzyme of D-amino acid stimulated ethylene production in *Arabidopsis thaliana*"

### Supplemental Figure S1

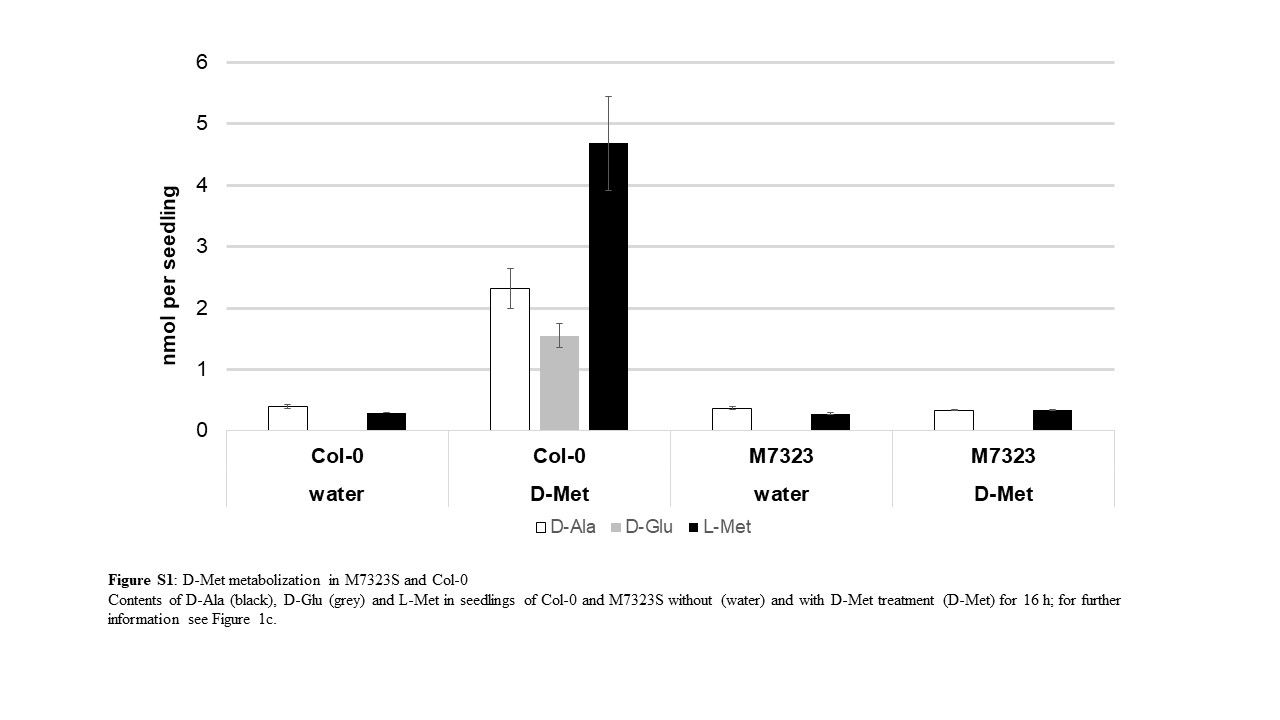

### Supplemental Figure S2a

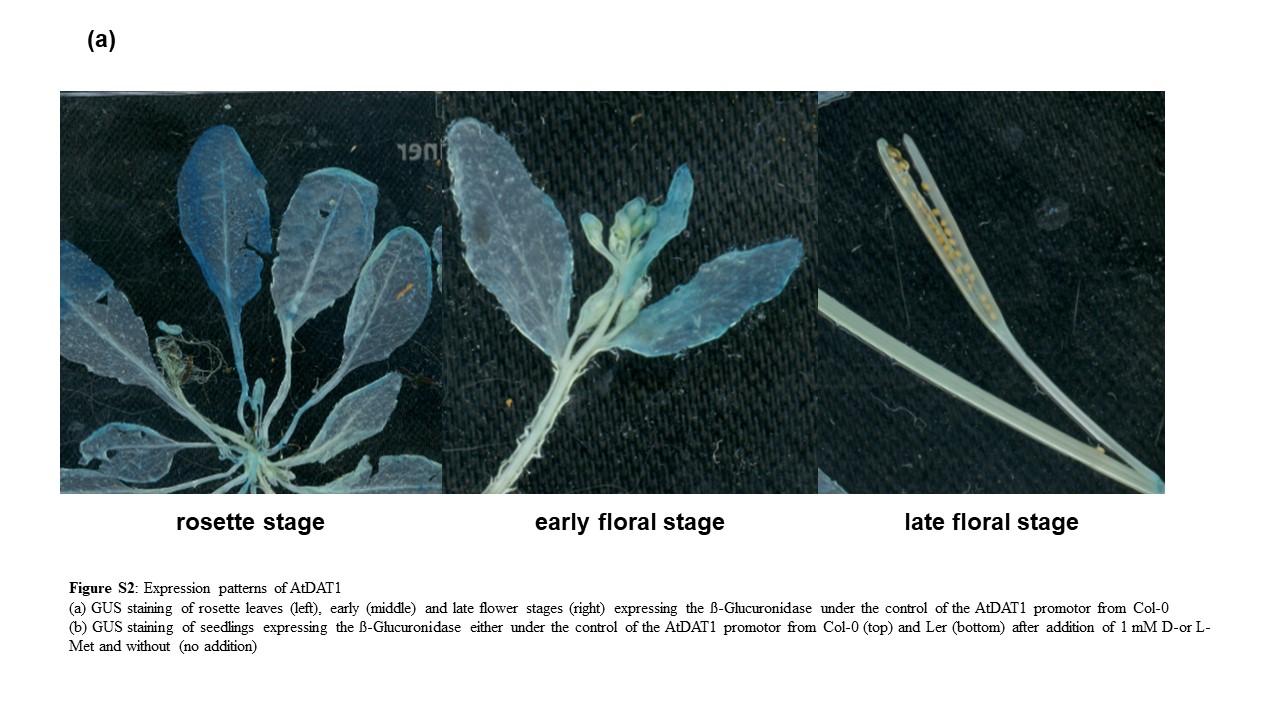

### Supplemental Figure S2b

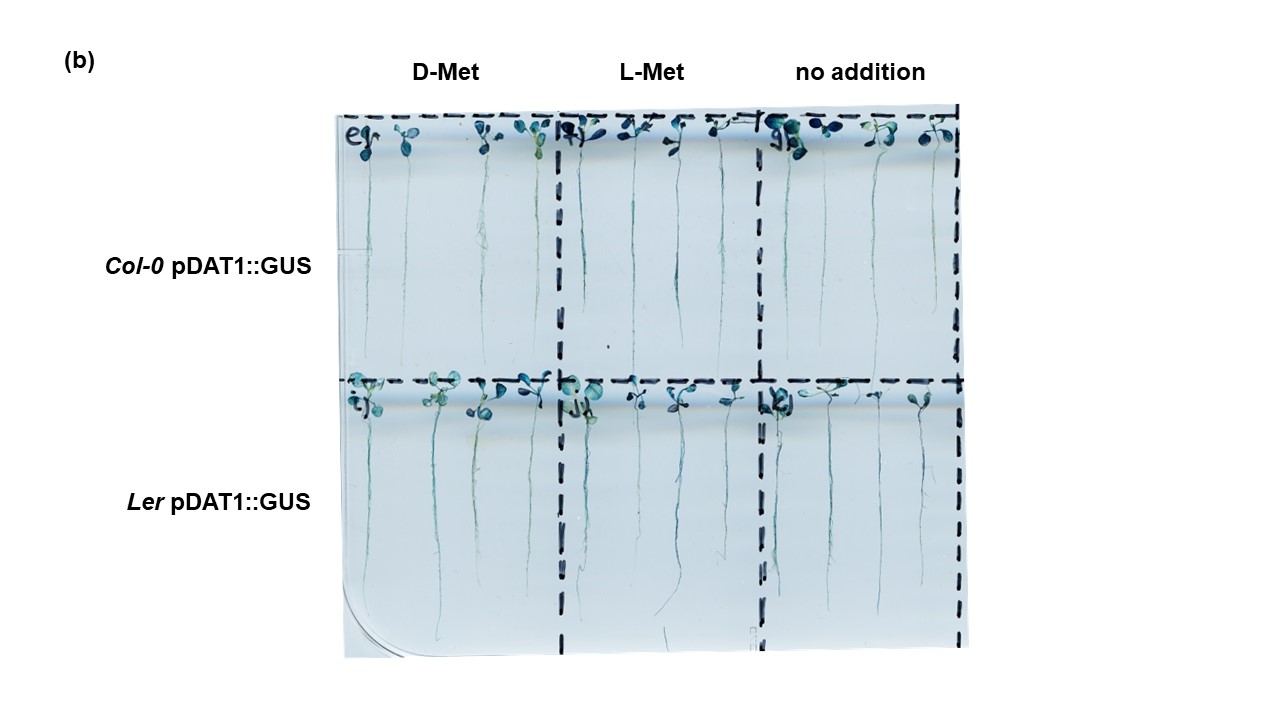

### Supplemental Figure S3

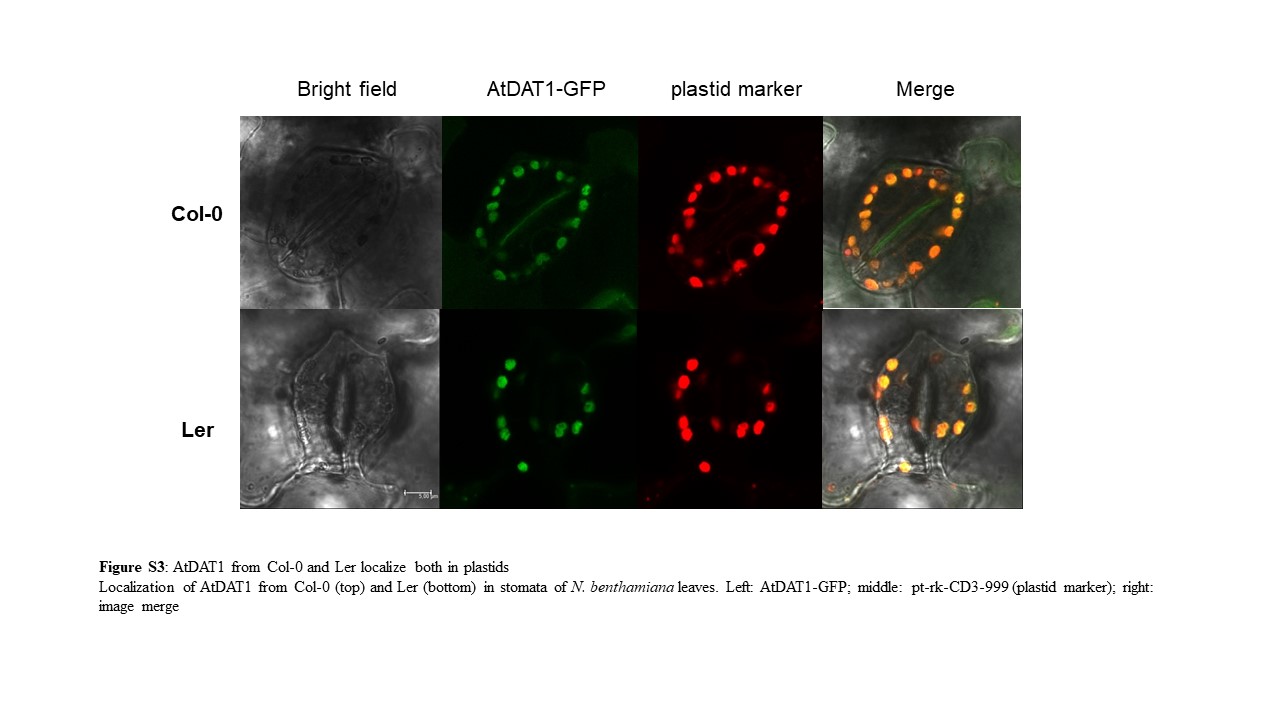

### Supplemental Figure S4

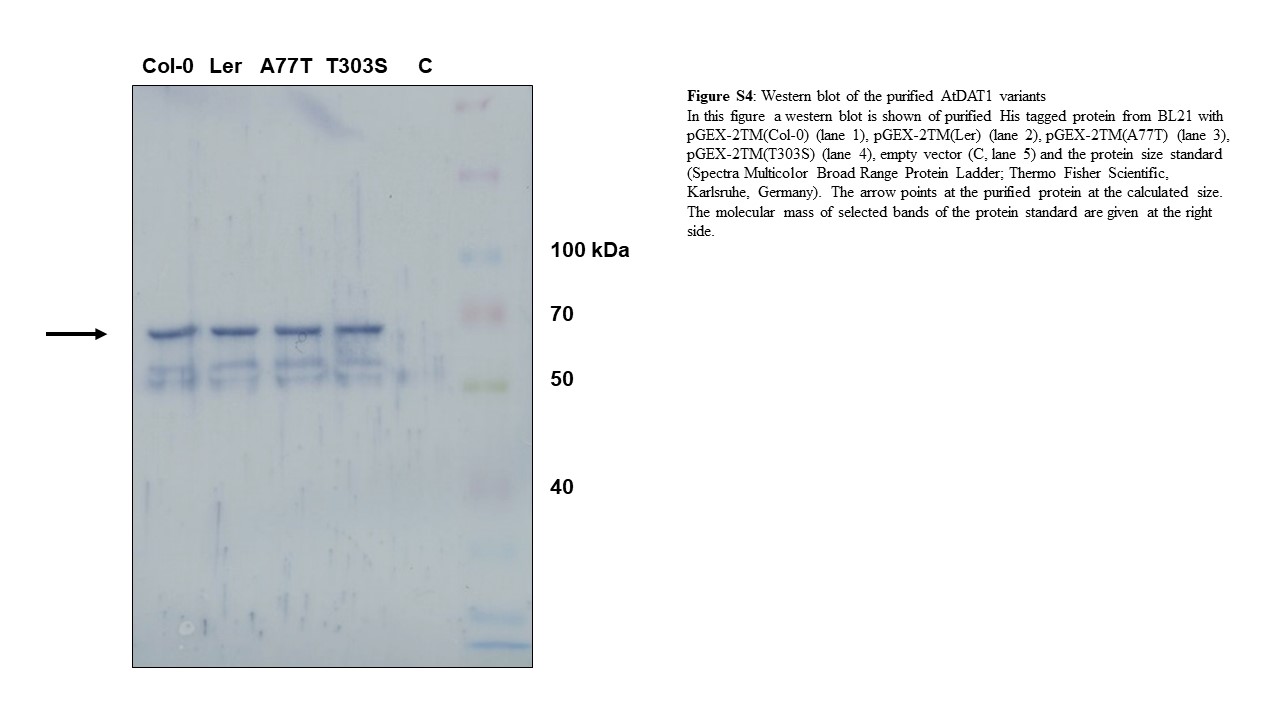

### Supplemental Figure S6

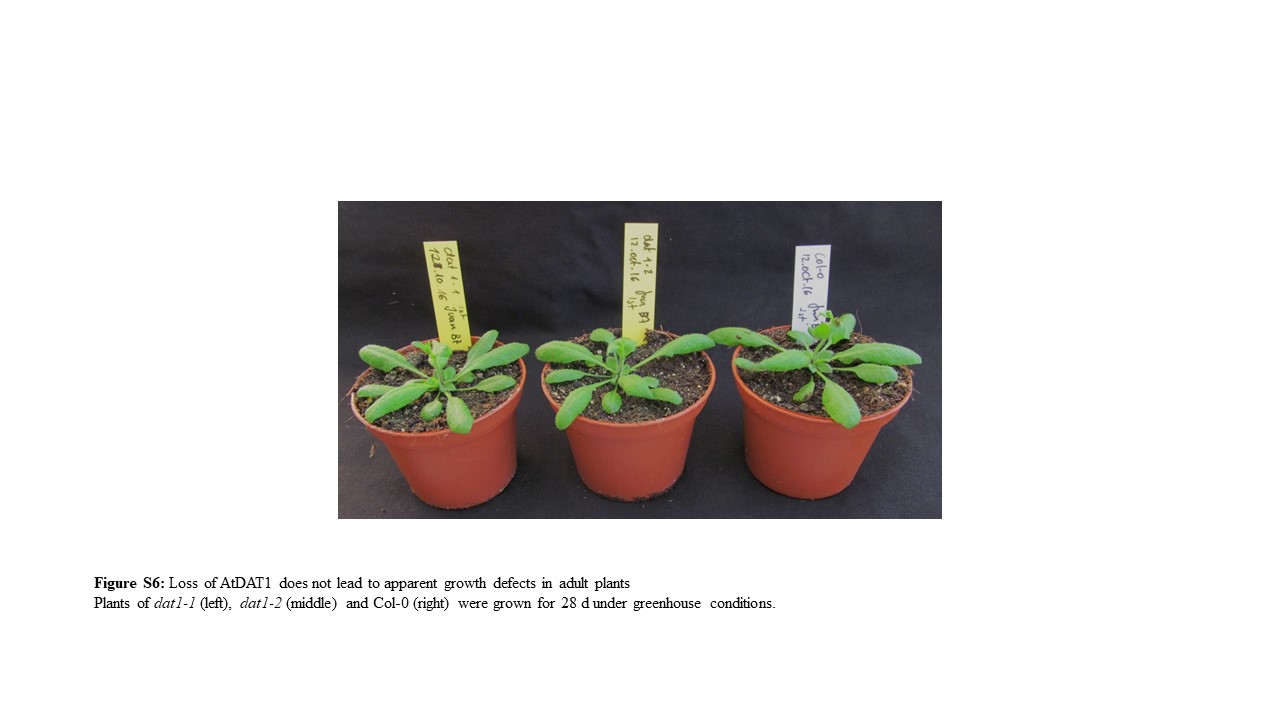

### Supplemental Figure S7

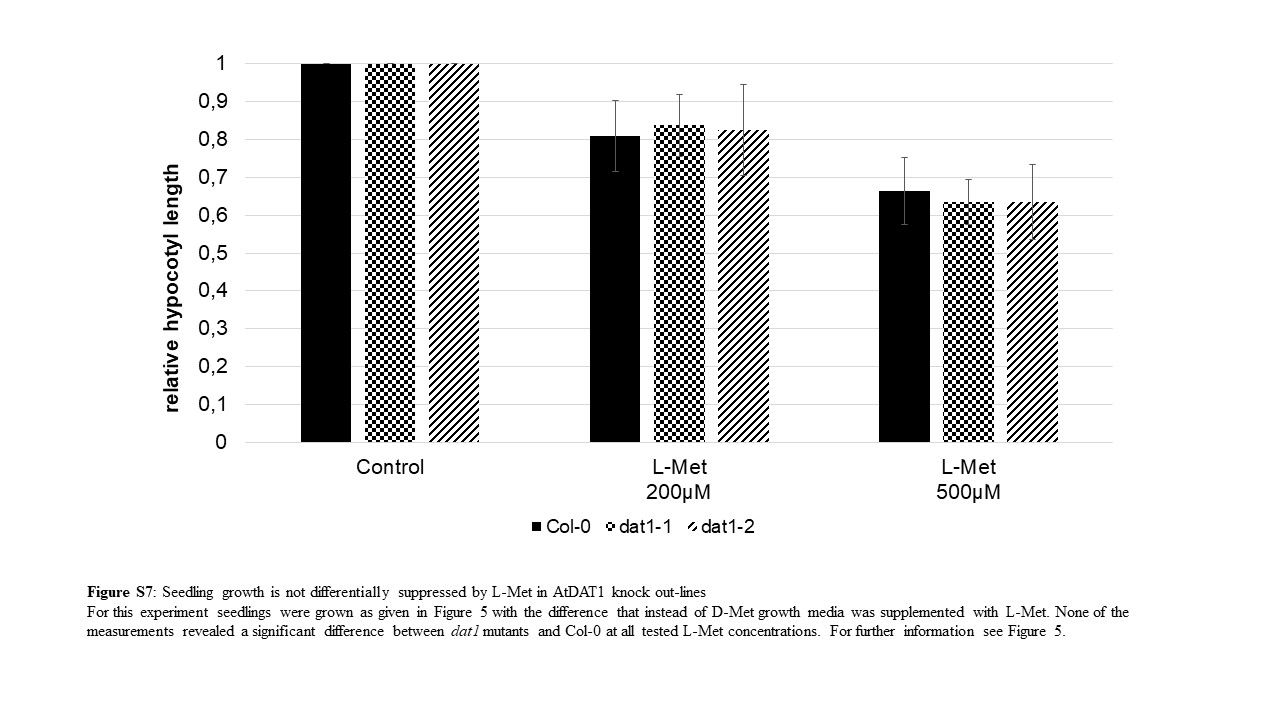
