## Supplemental Figure S5 for "AtDAT1 is a key enzyme of D-amino acid stimulated ethylene production in *Arabidopsis thaliana*"

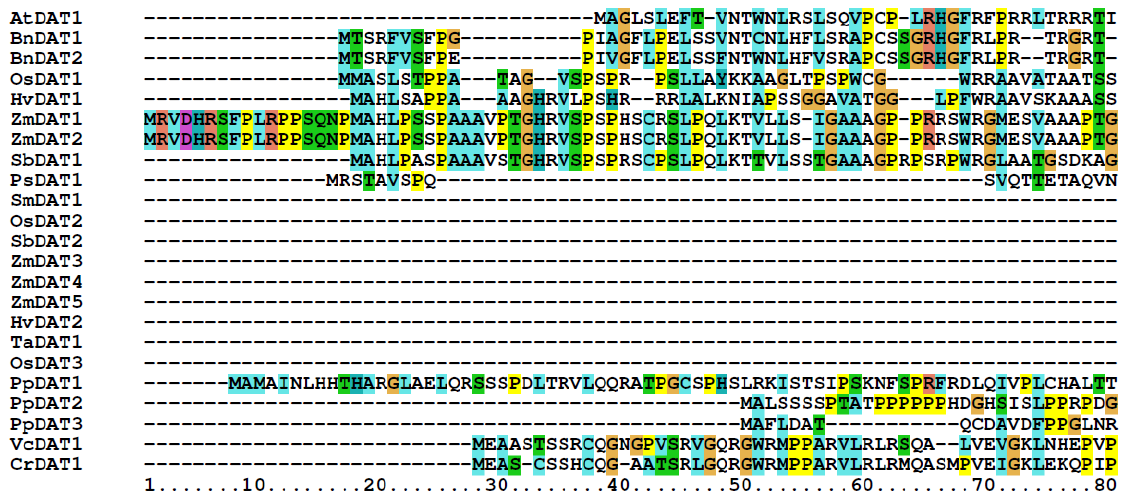

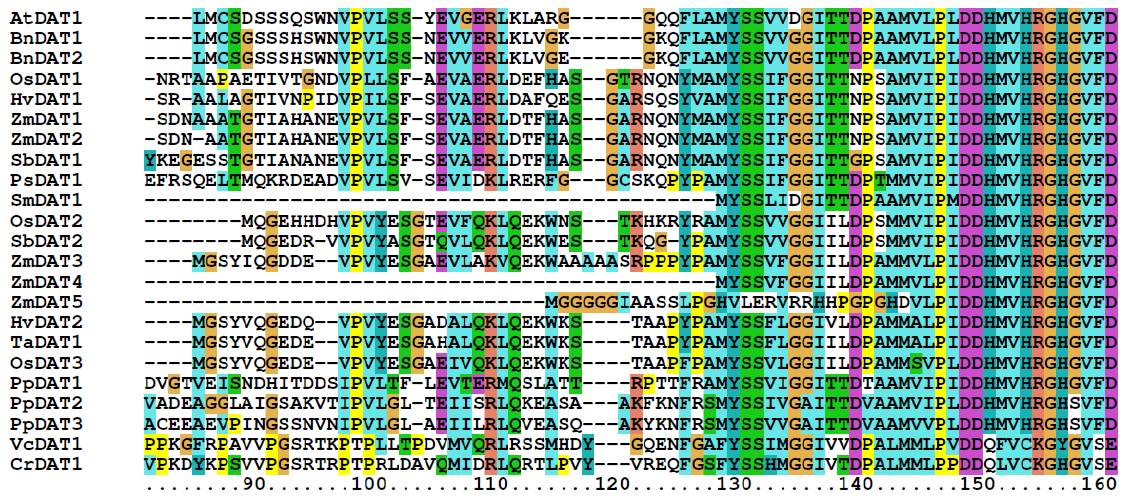

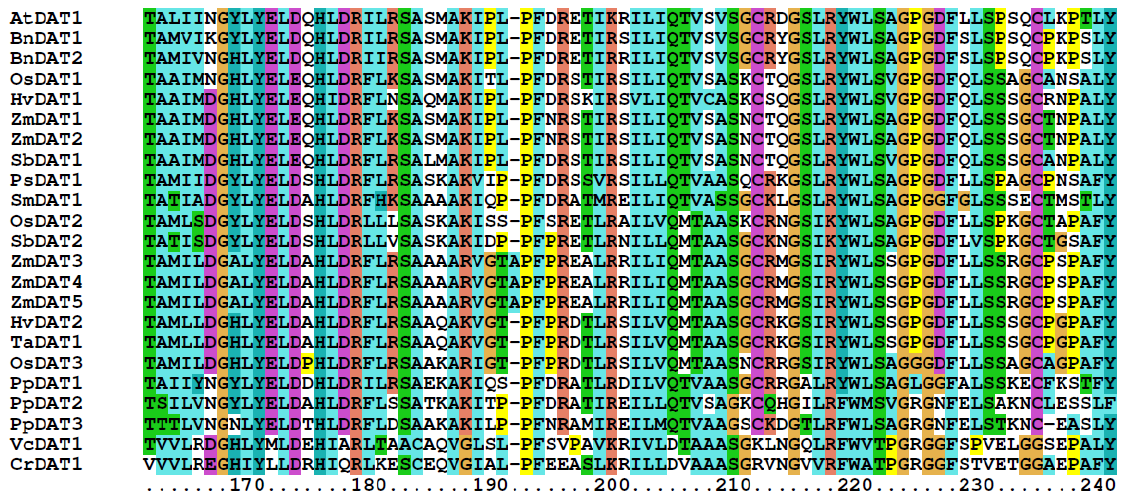

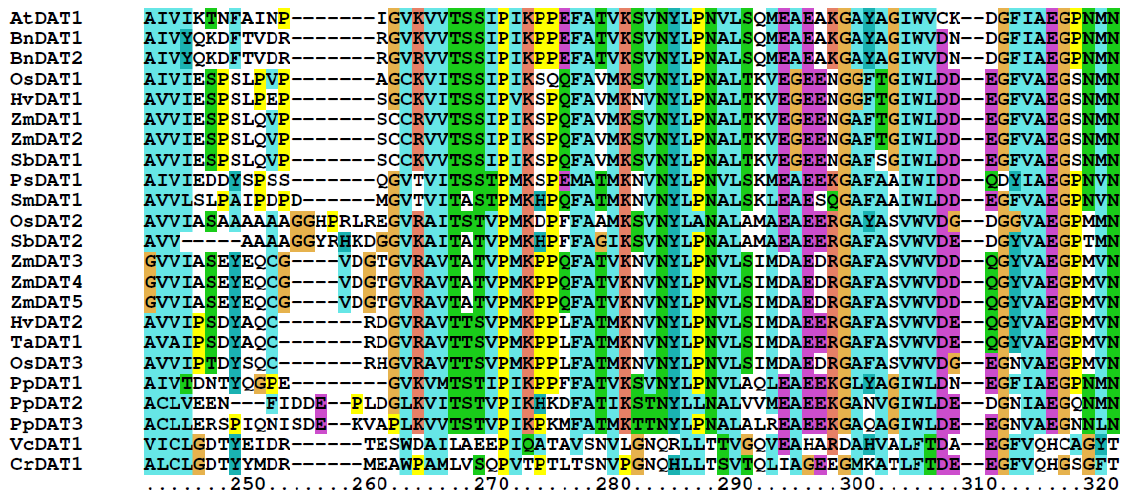

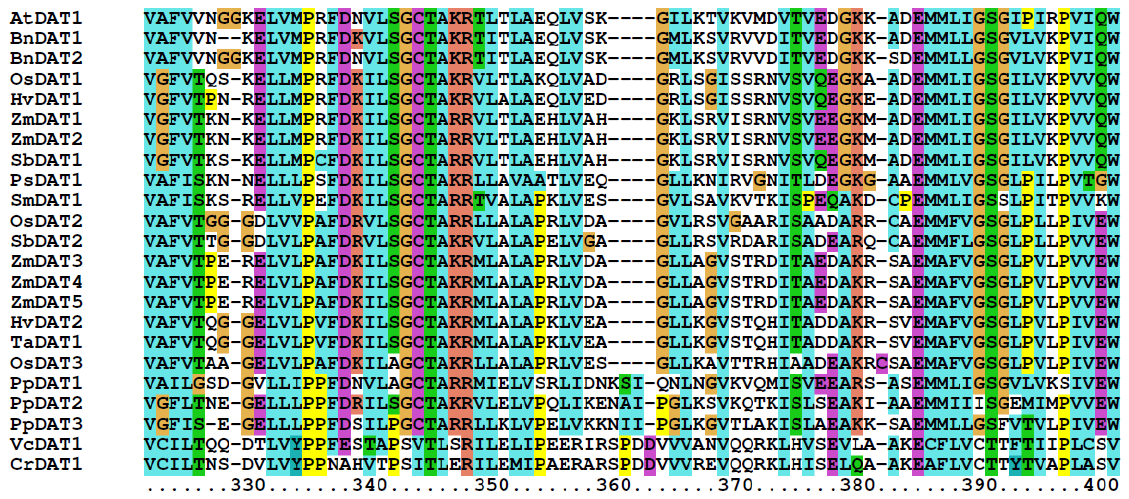

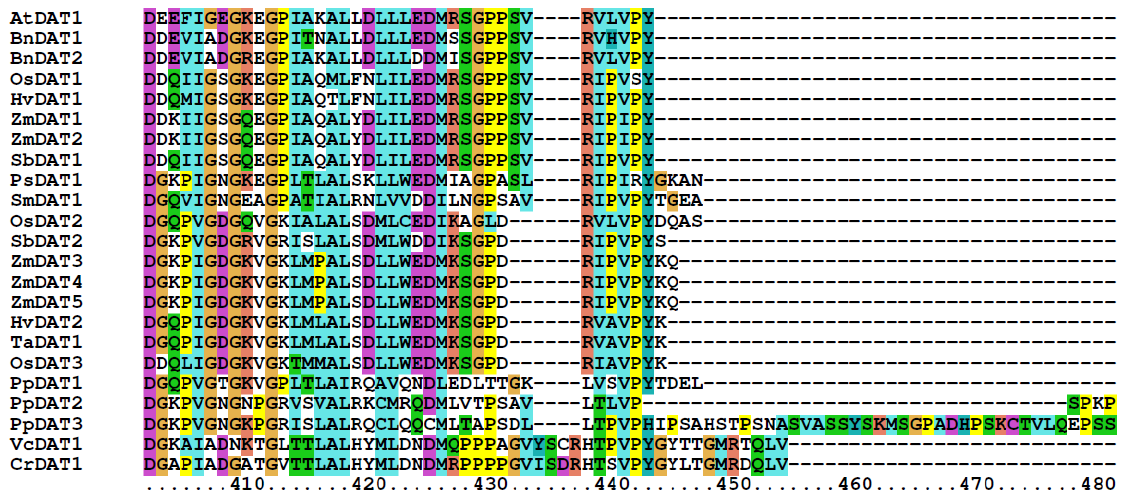

**Figure S5:** Alignment of DAT1 protein sequences from different plants and algae
Protein sequences were taken from the Phytozome 12 database (https://phytozome.jgi.doe.gov/pz/portal.html#) with AtDAT1 as search sequence. The alignment was constructed with ClustalX 2.1 (Larkin et al. 2007). AtDAT1 in the first line of the alignment is marked by a blue box. The chloroplastic transit peptide cleavage site (arrow) was determined according to the respective entry in the Plant Proteome Database (PPDB; http://ppdb.tc.cornell.edu/). Red triangles mark the sites of amino acid exchanges A77T and T303S between AtDAT1 from Col-0 and Ler. The blue star denotes the site of nonsense mutation in M7323S.
