## Supplemental Tables S1-3 for "AtDAT1 is a key enzyme of D-amino acid stimulated ethylene production in *Arabidopsis thaliana*"

Supplementary Tables

**Table S1** Primers used in this study

| **Primer name** | **Primer sequence** | **Purpose** |
| --- | --- | --- |
| DAT1-S1 | 5´-AGGTCTCTCTCTCAAGTTCCATGTC-3´ | genotyping |
| DAT1-Start | 5´-caccATGGCAGGTTTGTCGCTGGAGTTTACAG-3´ | cloning cDNA |
| DAT1-A1 | 5´-GTAAGGAACAAGAACACGAACGGAAG-3´ | genotyping, cloning cDNA |
| DAT1-RTS1 | 5´-CTCTATTCTTTTGAGTCCCAAACC-3´ | RT-PCR |
| DAT1-RTA2 | 5´-AGACAGATACCTAAAAACCCCATG-3´ | RT-PCR |
| ProDAT1-SGW | 5´-CACCCTCTTCTTCCGCTGCCGATTCACAAGAC-3´ | cloning DAT1-Promoter |
| ProDAT1-AGW | 5´-CAAACCTGCCATGGGGTTTGGGACT-3´ | cloning DAT1-Promoter |
| ACT2-F | 5´-TCCAAGCTGTTCTCTCCTTG-3´ | RT-PCR |
| ACT2-R | 5´-GAGGGCTGGAACAAGACTTC-3´ | RT-PCR |
| SALK-LB1 | 5´-AATCAGCTGTTGCCCGTCTCACTGGTGAA-3´ | genotyping |

**Table S2**: Comparison of the transaminase activity of AtDAT1 with different amino donors and acceptors
The transaminase activity (V_max_) of the *Col-0* variant of AtDAT1 was compared with D-Met, D-Trp and D-Ala as amino donor and 2-OG and pyruvate as acceptor molecules (in nmol product mg^-1^ Protein min^-1^); Each value represents the average of three independent assays (standard deviations are given in parentheses).

|  |  | **Amino donor** | | |
| --- | --- | --- | --- | --- |
|  |  | D-Met | D-Trp | D-Ala |
| **Amino acceptor** | Pyruvate | 2686.4 (±36.7) | 543.0 (±29.2) | Not determined |
|  | 2-OG | 16.3 (±0.8) | 5.1 (±0.7) | 17.1 (±2.7) |

**Table S3**: Relative malonyl-methionine contents (in %) in seedlings grown with or without addition of 10µM ACC
For these measurements seedlings of Col-0 and *dat1-1* were grown and analyzed as given in Figure 7.

|  | *-ACC* | *±SD* | *+ACC* | *±SD* |
| --- | --- | --- | --- | --- |
| Col-0 (light), 0 mM D-Met | 0.0 | 0.0 | 0.0 | 0.0 |
| Col-0 (light), 200 mM D-Met | 70.5 | 5.0 | 54.6 | 5.9 |
| Col-0 (light), 500 mM D-Met | 100.0 | 8.0 | 100.0 | 7.9 |
| *dat1-1* (light), 0 mM D-Met | 0.1 | 0.0 | 0.0 | 0.0 |
| *dat1-1* (light), 200 mM D-Met | 182.2 | 5.2 | 129.1 | 13.5 |
| *dat1-1* (light), 500 mM D-Met | 463.1 | 15.3 | 392.0 | 15.0 |
| Col-0 (dark), 0 mM D-Met | 0.0 | 0.0 | 0.0 | 0.0 |
| Col-0 (dark), 200 mM D-Met | 84.5 | 0.7 | 83.2 | 5.8 |
| Col-0 (dark), 500 mM D-Met | 100.0 | 2.8 | 100.0 | 8.7 |
| *dat1-1* (dark), 0 mM D-Met | 0.0 | 0.0 | 0.0 | 0.0 |
| *dat1-1* (dark), 200 mM D-Met | 128.5 | 9.8 | 95.6 | 2.7 |
| *dat1-1* (dark), 500 mM D-Met | 335.0 | 6.5 | 229.8 | 4.7 |
